# Heterogeneous pseudobulk simulation enables realistic benchmarking of cell-type deconvolution methods

**DOI:** 10.1101/2023.01.05.522919

**Authors:** Mengying Hu, Maria Chikina

## Abstract

Computational cell type deconvolution enables estimation of cell type abundance from bulk tissues and is important for understanding cell-cell interactions, especially in tumor tissues. With rapid development of deconvolution methods, many benchmarking studies have been published aiming for a comprehensive evaluation for these methods. Benchmarking studies rely on cell-type resolved single-cell RNA-seq data to create simulated pseudbulk datasets by adding individual cells-types in controlled proportions. In our work we show that the standard application of this approach, which uses randomly selected single cells, regardless of the intrinsic difference between them, generates synthetic bulk expression values that lack appropriate biological variance. We demonstrate why and how the current bulk simulation pipeline with random cells is unrealistic and propose a heterogeneous simulation strategy as a solution. Our heterogeneously simulated samples show realistic variance across hallmark gene-sets when comparing with real bulk samples from the TCGA dataset of the same tumor type. Using this new simulation pipeline to benchmark deconvolution methods we show that introducing biological heterogeneity has a notable effect on the results. Evaluating the robustness of different deconvolution approaches to heterogeneous simulation we find that reference-free methods that rely on simplex estimation perform poorly, marker-based methods and BayesPrism are most robust, while regress-based approaches fall in between. Importantly, we find that under the heterogeneous scenario marker based methods and BayesPrism outperform state of the art reference methods. Our findings highlight how different conceptual approaches can negate unmodeled heterogeneity and suggest that there is room for further methodological development.

## 1 Introduction

Bulk RNA-sequencing experiments reveal average gene expression values for all cells present in a sample mixture. Computational deconvolution methods separate the mixed signals from the aggregated expression and provide estimation of cellular components without physical isolations. The inferred cellular proportions are important to understand the ecosystem of the tissue and can be used as covariates in differential expression, reducing false positives and false negatives [1, 2]. Moreover, for heterogeneous bulk samples like tumor, deconvolution enables identification and quantification of the infiltrating immune populations, which provides rich prognostic values and can guide targeted therapy (e.g., in immunotherapy) [3, 4, 5, 6, 7].

Numerous deconvolution methodologies have been developed (see [1] for review), aiming at estimation of cell-type abundance from bulk transcriptomic data. Depending on a priori knowledge used, these methods can be broadly classified into two categories: reference free and reference based. Reference-free methods [8, 9, 10] are completely unsupervised and do not require any prior knowledge as input. Such methods are based on finding a simplex, which is a geometric data structure expected under ideal mixture proportion scenarios. Reference-based methods require cell type-specific profiles and depending on the underlying algorithm, they can be further classified into another three categories: regression-based, marker-based and Bayesian methods. Regression-based methods require an expression matrix as input, which consists of a cell type-specific expression profile for selected genes that can discriminate between cell types. These methods then solve the deconvolution as a regression problem. A comprehensive evaluation of factors involved in regression-based methods, like data transformation, normalization, and regression algorithms, can be found elsewhere [11]. Marker-based methods require a set of genes that characterize the expression patterns in different cell types and return either an enrichment score [12] that is unitless or abundance estimates [13, 14]. Bayesian method [15] uses an scRNA-seq reference as prior information and infers a joint posterior distribution of the cell type proportions and gene expression at the same time.

Rapid development of deconvolution methodologies now raises another challenge of evaluating their performance across diverse realistic settings. Many benchmarking studies have undertaken comprehensive evaluation of deconvolution methods under various testing conditions [2, 16, 11, 17]. Regardless of the focus of their evaluation, all benchmarking efforts rely on datasets with known ground truth. In the past ground truth was obtained using empirical approaches like fluorescence-activated cell sorting (FACS) or immunohistochemistry (IHC) staining of the same bulk RNA-seq samples [17]. However, these approaches are resource intensive resulting in datasets with limited sample size. An alternative approach is computational mixing where purified expressions of different cell-types are mixed in controlled proportions [12, 18]. While the purely computational strategy can generate large datasets this approach has the clear limitation that it makes the strong assumption that proportion variation and random noise are the only source of variance in the data.

Increasing availability of single-cell data offers the opportunity to create more realistic simulations. Instead of computational mixing of pure expression states, individual single cell profiles are added together in controlled proportions [11, 17, 19]. This has the explicit advantage over pure computational mixing as it introduces more variations in the simulated samples. However, as we will show in this work, while this approach has rapidly become the standard method for benchmarking deconvolution methods, the problem with unrealistic biological variance is only partially resolved. To simulate data compatible with bulk measurements, a large number of cells (typically hundreds) are added for each simulated sample. As such the pure cell expression in each sample, while not exactly identical, tends towards the global mean, enforcing the unrealistic assumption that there is no systematic variation beyond cell-type proportions. One possible solution is to take into account tumor heterogeneity in synthetic bulk mixtures. In Chu et al.’s study [15], they created such simulated bulk mixtures by restricting the malignant cells to aggregate for each simulated sample. However, the approach was not extended to other cell-types, and more importantly there is currently no general evaluation of how heterogeneity affects the deconvolution results.

In this study, we demonstrated why and how the current bulk simulation pipeline using random single cells is problematic and proposed a more realistic simulation strategy called heterogeneous simulation that can capture proper biological variance. Heterogeneously simulated bulk samples overcome the low-heterogeneity issue with current simulation pipeline and show high coordination with true bulk samples. We then provide an in-depth comparison between deconvolution methods from two major categories under different bulk simulation strategies using our curated benchmarking frameworks [20]. By summarizing deconvolution performance across experimental repeats, we found that introducing biological heterogeneity has a notable effect on the deconvolution results, with reference free methods being most affected, marker-based methods and Bayesian more robust, and regression-based methods in between. Our study can guide researchers in choosing the most appropriate deconvolution methods and proposes a highly realistic simulation framework that can guide further methodological development.

## 2 Results

### 2.1 Bulk simulation using random cells ignores heterogeneity within constituent cell-types

Heterogeneity of tumors between different patients with the same tumor type has long been recognized [21]. Despite similar histological appearance, different patients can have intrinsically different genomic landscapes. In clinical practice, this heterogeneity motivates molecular subtyping and enables personalized treatment protocols [22, 23]. We first visualized this intra-tumor heterogeneity using tSNE plot of malignant cells from a single-cell RNA seq medulloblastoma (MB) cohort [24] (Fig 1a). Medulloblastoma is a well-recognized heterogeneous brain cancer with four distinct subtypes based on genetic characteristics. WNT, SHH, Group 3 and Group 4 [25, 26]. According to the tSNE clustering, malignant cells within the same molecular subtypes are largely clustered together, indicating underlying differences between subtypes. Within each subgroup, further heterogeneity is identified for malignant cells according to their patient ID. Visualization of malignant cells from additional single-cell RNA seq cohorts [27, 28, 29] also confirmed this intra-heterogeneity pattern (Fig S1).

**Figure 1:**
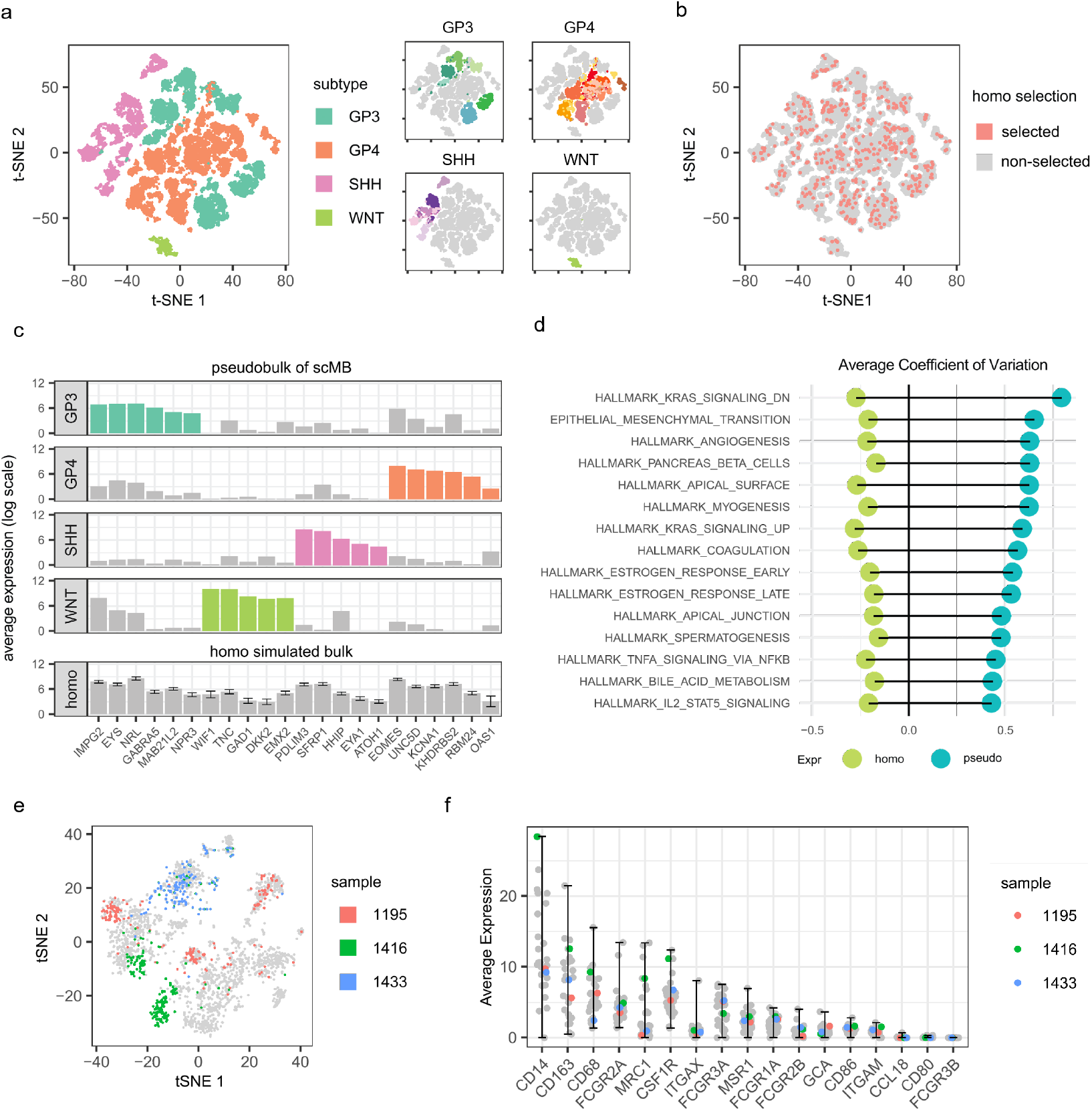
Homogeneous bulk simulation failed to retain intra-tumor variations. (a) tSNE plot of n=31,823 malignant cells from 29 medulloblastoma patients, colored by MB subtypes (left) and patient ID (right). (b) tSNE plot showing that 500 randomly selected malignant cells are scattered evenly within 31,823 malignant cells. (c) upper panel: average expression values of 22 MB subtype-specific genes from different subtypes of medulloblastoma patients reveals distinct patterns; lower panel: average expression values of 22 MB subtype-specific genes in 100 homogeneous simulated bulk samples, with errorbar indicating the 10th and 90th quantiles. (d) Hallmark expression variance comparison between 100 homogeneous simulated bulk samples and pseudobulk samples from 29 medulloblastoma patients. (e) tSNE plot of n=2,749 macrophage cells from the scMB dataset showing patient-specific heterogeneity in three randomly selected patients. (f) Average expression value of macrophage markers across macrophages in each sample, with error-bars representing the maximum and the minimum level, with three patients from (e) highlighted in colors.

Despite this general intra-tumor heterogeneity, random selection of cells from a heterogeneous population results in an evenly distributed selection of cells (Fig 1b), and such selection, if performed repeatedly, will create a homogeneous expression profile with low variance. To show this, we simulated 100 MB bulk samples using random cells (Methods: homogeneous simulation) derived from the single-cell MB cohort. We visualized the distribution of 22 MB subtyping specific genes [26] in the homogeneously simulated samples and compared them with the actual pseudobulk samples of the same single-cell cohort. We note that for the purpose of our work “pseudobulk” will refer to aggregation of cells from the same biological sample while datasets derived by combining cells across samples will be referred to as “simulated”. According to this distinction, pseudobulk samples should be very close to realistic bulk samples retaining all biological variation, whereas simulated pseudobulk may deviate considerably from the real distribution. Indeed, considering pseudobulk samples we find a distinct expression pattern of subtyping specific genes in different subtypes of patients, while in the simulated samples, there’s minimal heterogeneity in the expression values between samples, with the 10% and 90% expression quantile fluctuate around the average level, which is contradictory to the subtype specific patterns we expected to see (Fig 1c).

Additional evidence of low heterogeneity comes from the comparison of variances for biological pathways (Fig. 1 d). In this comparison, by averaging the gene-level variations for different set of hallmark genes, we find that the variance in simulated gene expression is much lower than in the pseudo-bulk samples (Fig. 1 d). This suggests that bulk simulation using random cells lacks realistic variance.

Moreover, since random cell selection is extended to cell types other than malignant cells in current bulk simulation practices, we asked whether the intra–heterogeneity pattern can be found in other cell types as well and answer the question whether random cell selection for non-malignant components is proper. We visualized the expression patterns for macrophages in the same single-cell cohort of MB using the tSNE plot (Fig. 1 e) and found that, similar to malignant populations, macrophage cells also tend to cluster within the same patients. Average expression values for macrophage markers [30] in different patients exhibited diverse expression patterns (Fig 1f), suggesting that even for some well-established cell-type specific markers, intra cell-type heterogeneity between patients is not uncommon. Additional evidence came from other immune cell-types and other single-cell cohorts, where we also observed an uneven distribution of expressions for immune cell specific genes (SupplementaryTable1) and a patient-specific clustering pattern within non-malignant cells (Fig S2-5). This suggests that by random combination of non-malignant cells, intra-heterogeneity is no longer preserved.

Together, we conclude that bulk simulation by combining random cells from a heterogeneous population ‘averages out’ the meaningful heterogeneity between cells and creates a homogeneous bulk profile that is unrealistic and lacks proper variance.

### 2.2 A heterogeneous bulk simulation strategy

Based on the observation of patient-specific heterogeneous expression patterns for different celltypes, we designed a novel bulk simulation approach that retains the heterogeneity between samples by restricting the cells to use when building the bulk mixtures (Fig 2). This means that instead of aggregating single cells randomly from the same cell-types, we restrict that each cellular component of a synthetic bulk sample is only made up of single cells from the same patient, while the combination of different cellular components is random (Methods). In this way, intra-heterogeneity within constituent cell-types is maintained while the bulk samples still have proper randomness. Theoretically, by random combinations of patient-specific cellular components, we can get *n^k^* numbers of combinations, where n is the number of patients in the single cell cohort and k is the number of cell-types identified. But in reality, some cell-types can be missing from one patient or only have a limited number of cells. Therefore we introduce a threshold parameter to specify the limited number of cells required to build each cell-type, and combine cells from different patients if the threshold is not met. This step creates extra randomness into the simulation pipeline and ensures that each simulated bulk sample contains proper cellular components.

**Figure 2:**
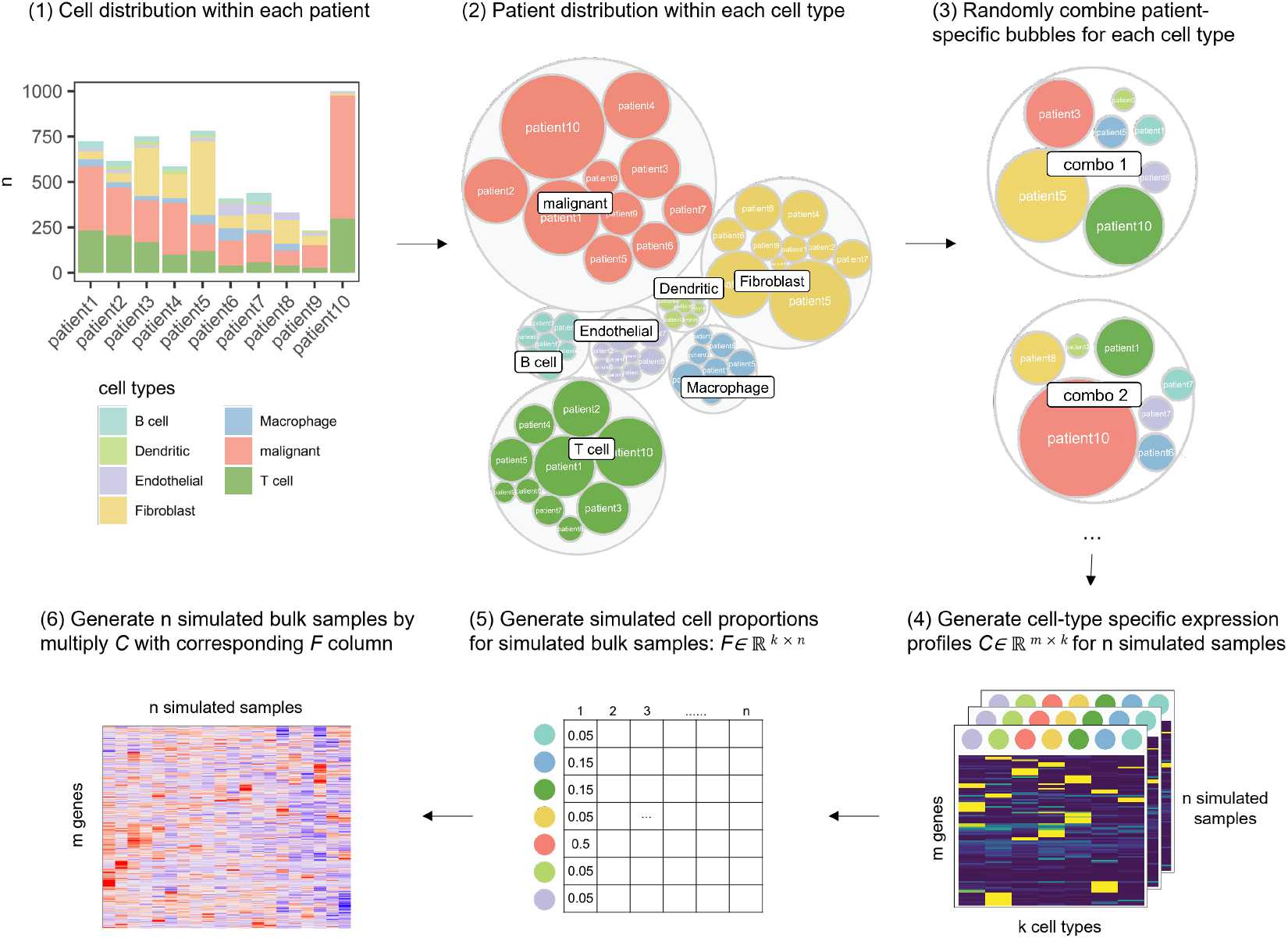
Schematic representation of heterogeneous bulk simulation. Heterogeneous bulk simulation involves the following steps: (1) cell-type distribution is first summarized at per patient level; (2) patient barcode is summarized at per cell-type level; (3) selecting single cells to aggregated for *n* simulated bulk samples by restricting each cell-type from the same patient and random combining cell groups; (4) generating cell-type specific expression profile *C* using the aggregated single cell profile from step(3); (5) generating simulated cell-type proportions F for simulated bulk samples; (6) cell-type specific expression profile C is weighted by cell-type proportions *F* to generate the final heterogeneously simulated bulk profiles.

As is done in previous benchmarking efforts in order to build a ground-truth dataset for deconvolution benchmarking [11], we first generate sample-level cell-proportion vectors and then add single cells in corresponding proportions with the added heterogeneity preserving restriction described above (Fig 2). Our method also differs from previous approaches in how the proportion vectors are generated. Previous practices of generating such predefined proportion profiles used either uniform distribution or normal distribution [11, 31, 19]. However, cellular proportions may not strictly fall into these distributions and we reasoned that beta distribution is more appropriate in capturing the true scenario. Using cellular fractions observed from single cell expression data as an approximation for the true cellular fractions in real bulk samples, we fitted a beta distribution for each cell type. To generate proportion vectors each cell-type is sampled from its own beta distribution and the results are normalized to sum to 1. We compared our simulated fraction with the ‘runif’ simulation strategy used in Cobos et al.’s [11] study and find that this procedure matches the proportion distributions of the original pseudobulk samples (Fig S8).

We also consider a variation of our heterogeneous simulation strategy called ‘semi-heterogeneous’ simulation is also included in our later benchmarking framework. The only difference is that instead of restricting the cells to use for all cell types, semi-heterogeneous simulation aggregates random cells for all non-malignant cells and only requires the malignant cells to be restricted to one patient at a time. Semi-heterogeneous simulation was previously implemented in Chu et al.’s study [15] to benchmark their methods against other deconvolution tools. We will show in later sections how semi-heterogeneous simulated bulk samples maintain an intermediate level of biological variance.

The effect of the heterogeneous strategy is to introduce a level of model-misspecification. First and foremost this simulation breaks the assumption that all of the variation across samples can be explained by cell-type proportions. From the perspective of regression the heterogeneous simulation can also be seen as introducing a mean misspecification as the real cell-type mean of an individual sample will be different from that encoded in the reference matrix. This situation also mirrors the realistic scenario where the reference data and the bulk data differ due to non-biological factors such as RNA processing and platform effects (e.g. ribo depletion, common for bulk, vs. poly-A selection, standard for single cell).

### 2.3 Comparison between different bulk simulation strategies

In the previous sections, we showed the underlying problems with homogeneous simulation using random cells and introduced heterogeneous simulation as an alternative. We also introduce the idea of ‘semi-heterogeneous’ which was previously implemented in another study. Next, we want to compare the simulated datasets generated under different strategies and find the ones that maximally resemble the real data. We used four publicly available single-cell datasets (i) head and neck squamous cell carcinomas (HNSCC) tumors from Puram et al. [29], (ii) melanoma tumors from Schelker et al. [28], (iii) medulloblastoma (MB) tumors from Riemondy et al. [24] and (iv) another melanoma cohort from Jerby-Arnon et al. [27] as source input (Table 1) and applied three simulation strategies to each of them. This resulted in 12 simulated datasets, each of which is made up of 100 simulated samples. To create baseline bulk expression to compare against, we aggregated single cells from the same patients into pseudobulks and used them as approximations of real bulk samples. We also collected expression profiles from the TCGA datasets [32] when the pairing tumors are available.

**Table 1:**
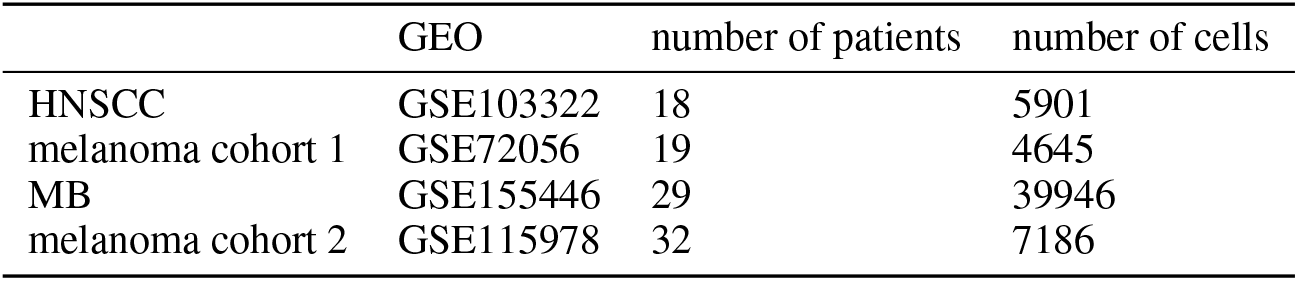
Single cell datasets used in this study

Distribution of mean, variance and coefficient of variation (CV) of gene expression levels are usually used to describe the characteristics of expression matrices [33], so we calculated these variables accordingly for both simulated datasets and real bulk datasets. By comparing gene-level CV across the samples in the simulated melanoma samples from GSE115978 dataset, we found that the heter-simulated bulk samples retain comparable variance with that of real bulk samples, while the homo-simulated bulk samples are in general less variable (Fig 3a) and the semi-simulated samples are in between. Summarized CV at pathway level [34] further confirmed this finding, and by extending the measurement of CV in the real TCGA bulk samples, we showed that heter-simulated samples retain proper biological variance compared with real bulk samples (Fig 3b). Additionally, by summarizing the pairwise correlation between simulated samples under different strategies, we found homo-simulated samples are highly correlated with each other, while heter-simulated samples have the least pairwise similarities (Fig. 3c, S6). A mean-variance plot further suggests that heter-simulated samples are comparable to real bulk samples (Fig 3d). A comprehensive variance comparison of hallmark pathways in 12 simulated datasets is shown in Fig 3e and S7, with all the heter-simulated samples resembling the actual variance in real data, and the homo-simulated samples containing the least variances. Of note, in the MB cohorts, the semi-simulated and heter-simulated samples show little difference with respect to pathway variances, this is because the simulated MB samples are mainly dominated by malignant cells, making it less distinguishable between two strategies.

**Figure 3:**
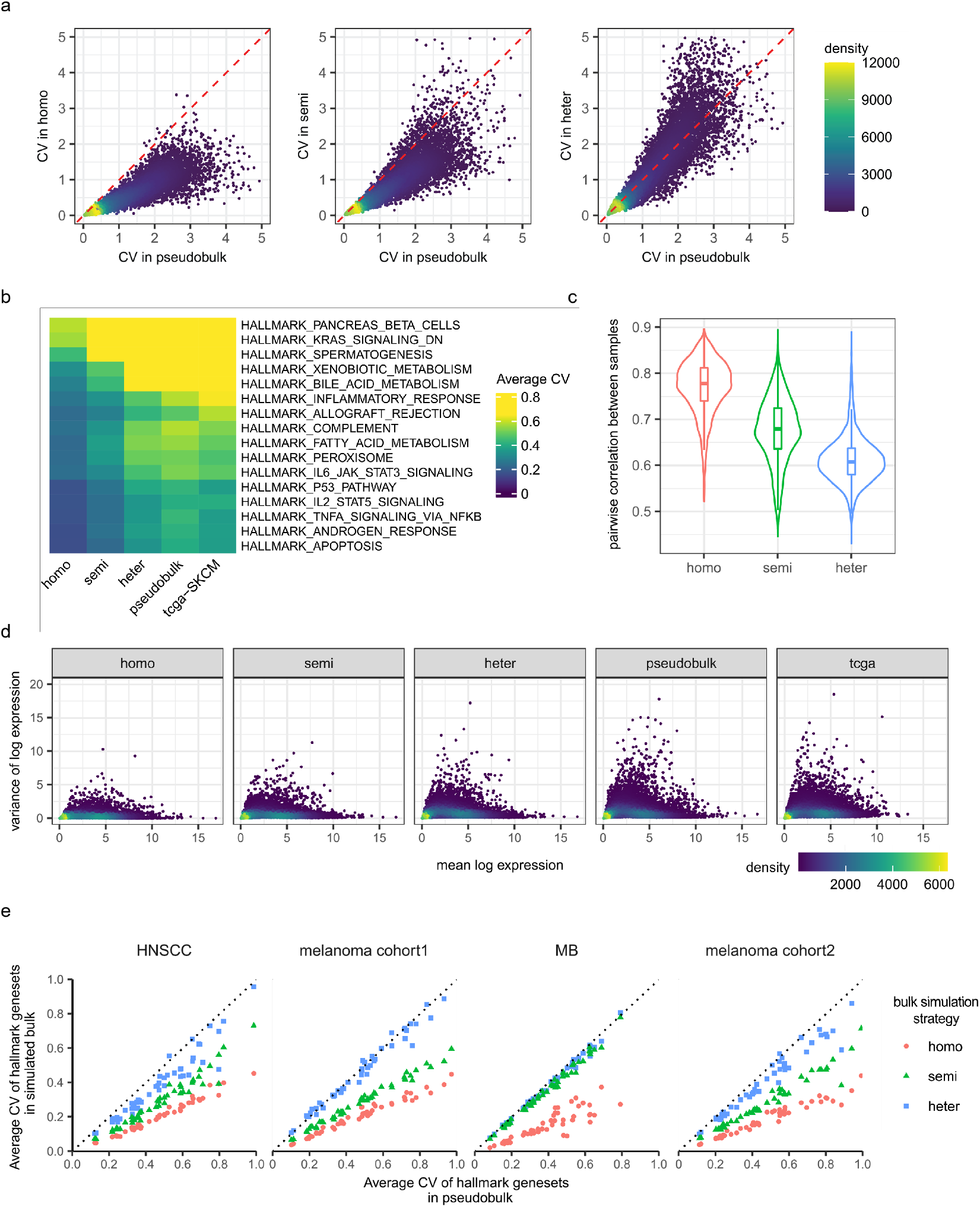
Heterogeneous bulk simulation maintains meaningful biological variations. (a) Scatter plots comparing coefficients of variation for all genes between simulated bulk samples and pseudobulk samples from 32 melanoma patients in single cell cohort GSE115978. From left to right, simulated bulk samples are generated through homo, semi-heter and heter simulation strategies, pseduobulk samples on the x axis are used as approximations to real bulk samples. (b) Heatmap showing average coefficients of variance for genes in 16 different hallmark gene-sets across five expression matrices: three simulated bulk expression matrices under different simulation strategies, one pseudobulk expression matrix from 32 melanoma patients and one real bulk expression profile from melanoma tumors in TCGA. (c) distribution of pairwise correlations between simulated bulk samples under different simulation strategies. (d) mean-variance plots for all genes in five expression matrices as in (b). (e) scatter plots comparing variance for all 50 hallmark gene-sets between the simulated samples and pseudobulk samples from four different single-cell cohorts, colored by simulation strategies.

Together, our results suggest that heterogeneous simulated bulk samples retain the characteristic of real bulk samples in terms of variability, biological variance, and distribution of expression levels. Stepping through homo, semi, and heter simulation, the heterogeneity level inside samples is increasing with the final heter being very similar to both pseudobulk and real bulk samples.

### 2.4 Assessment of deconvolution performance under different bulk simulation strategies

We designed a benchmarking framework that is adapted from Cobos et al. [11] (Fig 4, Method). Briefly, the framework takes a single cell expression profile as input and examines the deconvolution performance on the simulated bulk samples reconstructed from the single cells, using either homogeneous, heterogeneous or semi-heterogeneous simulation strategies as mentioned above. We included more than 8 deconvolution methods belonging to different categories of deconvolution methodologies. For reference-free methods, we selected CAMTHC [14] and linseed [9]. For regression-based methods, we generated the reference matrix using CIBERSORTx [35], autogeneS [36] and differential expression analysis, and solved the deconvolution problem using non-negative least squares regression (nnls). For marker-based methods, we implemented CAMTHC-marker [14] and gsva [37] using the same DE-genes that were used to build the reference matrix, and xCell [12] that contains built-in gene-lists. We also investigated a recently published method Bayesian method, BayesPrism [15]. We consider this method to be in its own class as while it makes use of a fully quantitative reference matrix it does so in a way that is most conceptually similar to a marker-based approach (see Discussion). We also incorporated two extra methods that deconvolute cell-type fractions using built-in reference matrices: epic [38] and quanTIseq [18]. The detailed mapping procedure from xCell scores, epic score and quanTiseq score to the matched cell-types can be found in Methods and SupplementaryTable2.

**Figure 4:**
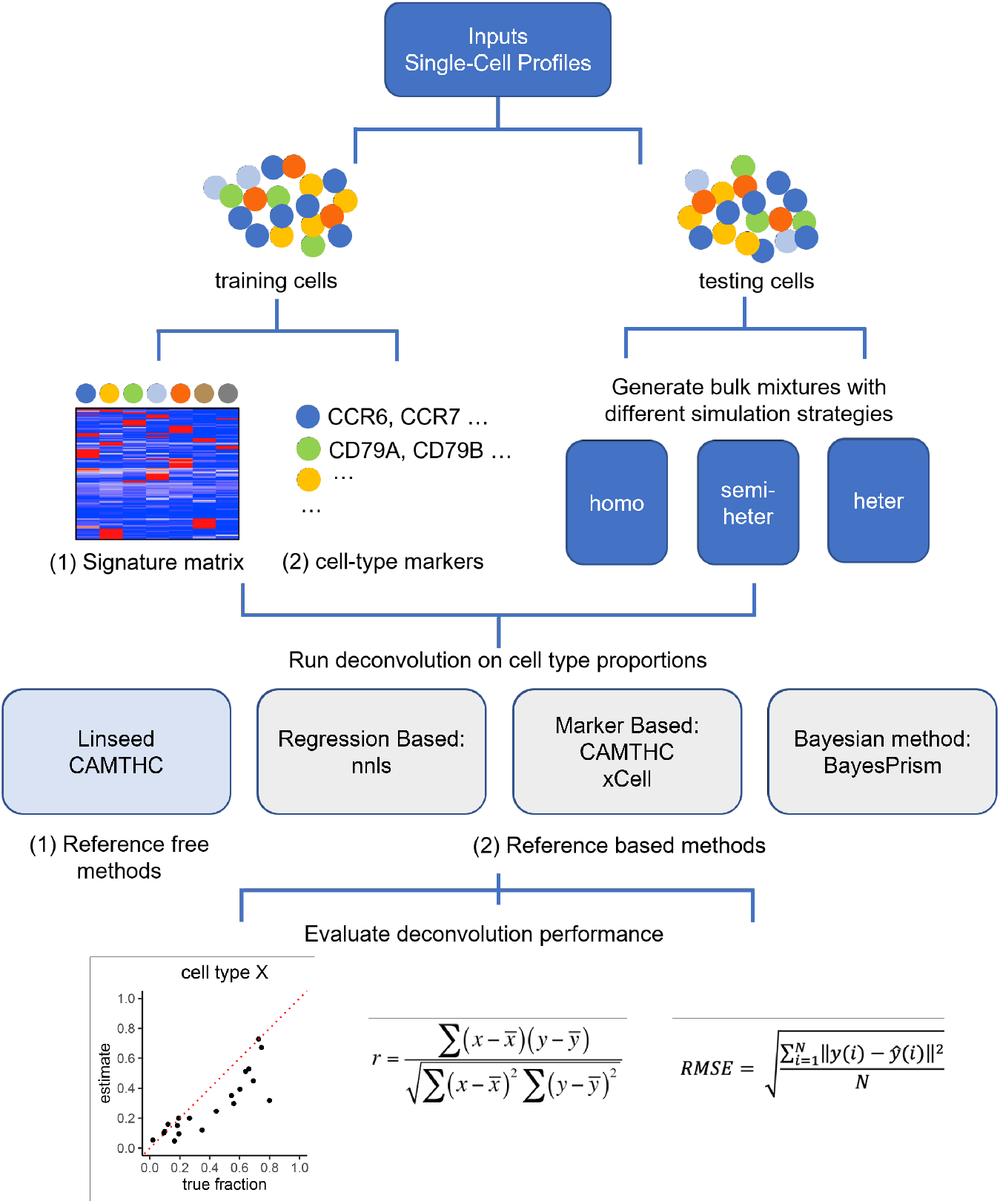
Schematic framework of the benchmarking study. The complete framework involves the following steps: (1) The benchmarking framework takes the single-cell expression matrix and predefined cell-type labels as input. (2) Single cell profiles are then splitted into training (50%) and testing (50%) sets. (3) Training set is used to create reference signature matrices consisting of informative marker gene expression, and cell-type marker list that can discriminate between different cell-types. Testing set is used to build synthetic bulk samples using three different simulation strategies. (4) Three categories of deconvolution methods are implemented: reference-based method, marker-based method and reference-free method. (5) Performance of different deconvolution methods is evaluated by comparing between the estimated and known fraction using the following statistics: Pearson correlation coefficient, root mean square error.

Procedures to evaluate deconvolution results vary in terms of whether the agreement between ground truth and inferred proportions is assessed by correlation or squared error and whether performance is evaluated per-cell type or globally. We focus our evaluation on per cell-type Pearson correlation, which reflects the accuracy of downstream inference such as the difference in proportions between two groups. We also evaluate root mean square error (RMSE) values, which evaluate if the inferred proportions are correct on the absolute scale across different cell types, with smaller RMSE indicating better performance. The deconvolution pipelines including simulation, deconvolution, and evaluation are then applied to four published single cell cohorts [24, 27, 28, 29] as mentioned in the previous sections and repeated 10 times for each cohort.

To understand how different bulk simulation strategies can affect the deconvolution results, we averaged cell-type level performance over 10 experimental repeats in terms of Pearson correlation and visualized them in boxplot. In Fig 5a, we mainly compared the performance of deconvolution methods under homo and heter simulation settings, with the formal representing bulk simulation in most current benchmarking work, and the latter representing real bulk samples with cross sample heterogeneity preserved. We showed that increasing heterogeneity results in a global performance drop for all the deconvolution methods. This is particularly striking for reference free methods that perform well under homogeneous simulation but poorly for the heterogeneous ones. Marker-based methods and the Bayesian method BayesPrism are less affected and have in general more consistent performance across datasets. For regression-based methods, using signature matrix generated from CIBERSORTx, autogeneS and DE analysis all showed good performance under homo setting, however, when bulk samples are heter-simulated, regression-based methods showed clear performance drop and are less consistent over experimental repeats.

**Figure 5:**
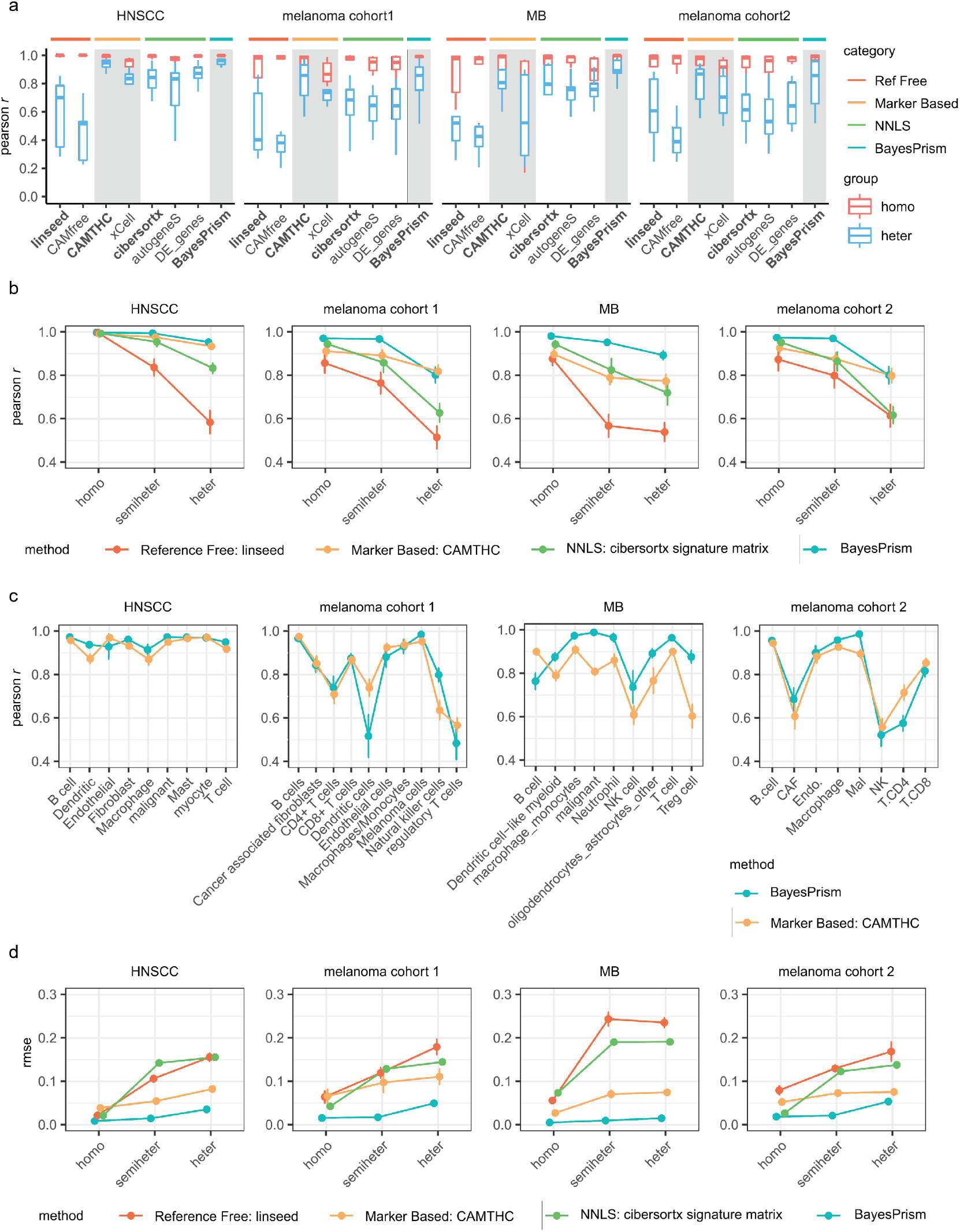
Impact of bulk simulation strategies on deconvolution performance. (a) Deconvolution performance, estimated by Pearson *r* (y axis) is summarized at per method level (x axis) for each single cell dataset used for benchmarking, with each box indicating the distribution of per cell-type level performance and colored by simulation strategies; deconvolution method with the best performance for each deconvolution category is highlighted in bold font. (2) Lineplot depicting the overall Pearson r (averaged over all cell-types) for four representing deconvolution methods across experimental repeats in four single cell datasets, with errorbar indicating the confidence intervals over experimental repeats. (3) Lineplot comparing cell-type level performance between CAMTHC and BayesPrism for heterogeneously simulated bulk samples, with errorbar indicating confidence intervals over experimental repeats. (4) Lineplot showing the distribution of global RMSE values across experimental repeats in four single cell datasets, with errorbar indicating the confidence intervals over experimental repeats.

A highly condensed comparison is shown in Fig 5b, where we picked up the best performed deconvolution methods from each category and summarized the cell-type level Pearson correlations into an overall performance score (Methods), with the error bar indicating the variance between experimental repeats. A key observation, similar to in Fig 5a, is that introducing heterogeneity into the bulk samples has a notable effect on the deconvolution performance, with marker based and Bayesian method being more robust, reference free method suffering the most and regression-based method lying between. Additionally, we found that Bayesian method ‘BayesPrism’ is consistently outperforming other methods under semi-heter simulation setting across the datasets, this is probably because under such setting, major heterogeneity between samples come from malignant cells, which is precisely captured by BayesPrism with its ‘cell.state.label’ option. As heterogeneity is not limited by malignant cells, for example, in two melanoma datasets and one HNSCC dataset which contain less malignant contents, BayesPrism shows no clear advantage over the marker-based method. Moreover, we observed that in the MB dataset performance drop from semi-heter to heter for deconvolution methods beyond BayesPrism is less obvious, this is probably due to the high malignant content in MB patients (with 90% of malignant cells on average), weakening the impact of patient-specific heterogeneity from the non-malignant cells.

We next focused on performance comparison for heterogeneously simulated bulk samples, which has been shown earlier to mostly resemble real bulk samples. For two deconvolution methods CAMTHC and BayesPrism that showed similar performance in Fig 5b, we extended the comparison to celltype level and found a similar result: for dataset with heterogeneity level spreaded between all cell types, for example in two melanoma datasets and one HNSCC dataset, BayesPrism showed similar performance compared to CAMTHC; however, when bulk samples are dominated by malignant content, BayesPrism outperformed CAMTHC in nearly all the cell types. We further reasoned that the prior information learned from training data can be affected when there is high malignant content, making the cell-type specific genes derived from DE analysis less reliable, which in turn weakens the deconvolution ability of CAMTHC.

Using global RMSE values as another performance measurement, we found a similar trend as we observed earlier, with RMSE values increasing as we introduce heterogeneity. BayesPrism achieves the lowest RMSE values under different simulation conditions, while CAMTHC ranked next. However, it should be noted that BayesPrism archives good performance at the expense of computational complexity. We compared the average running time required for CAMTHC and BayesPrism (Fig S14) and showed that BayesPrism is much more computational intensive, while CAMTHC resolves the deconvolution problem only in seconds, making CAMTHC a more effective option given the same input type. Detailed performance results of deconvolution methods, with regard to per-cell-type Pearson *r* and global RMSE values for four simulated datasets can be found in Fig S9-S13.

Taken together, our results highlight the effect of sample-specific heterogeneity in deconvolution results and suggest that previous benchmarking work using homogeneous samples does not reflect real world performance. We show that while reference/regression based methods are most widely used these are not top performers in our analysis. Instead we show that BayesPrism and CAMTHC, neither of which is regression based, achieve the best deconvolution performance under realistic bulk simulation settings.

## 3 Discussion

In this study, we introduced the importance of heterogeneity in bulk sample simulation and examined how heterogeneity could influence the deconvolution results. We investigated two major categories of deconvolution methods by applying them to simulated bulk samples with different heterogeneity levels and identified the top performing ones in each category. Our results showed that marker-based and BayesPrism are less affected by heterogeneity, while reference-free methods are most affected and reference-based methods lying in between.

One of our key findings is that reference free methods perform poorly under the heterogeneous scenario. Reference free methods are attractive as they require no prior knowledge and it has been repeatedly suggested that these methods produce reliable proportion estimates [8, 9]. However, we show that the more realistic simulation strategy the accuracy is much lower than would be expected from the previously reported results. Reference free methods rely on fitting a simplex structure which is the expected data geometry if the only source of variation is cell-type proportions. However, adding variation beyond cell-type proportions introduces additional lower dimensional structure making the proportion associated simplex structure more difficult to isolate.

We also report that both BayesPrism and marker-based methods are robust to heterogeneity and in particular can outperform reference based approaches. Our results align with a recent study [39] that benchmarked deconvolution methods on real bulk and single cell data finding that BayesPrism strongly outperforms all tested reference based methods when evaluated for consistency across different biochemical and bioinformatic processing pipelines for the same biological sample. While the Hippen et al’s study looks at heterogeneity from a different perspective, the fundamental conclusion that reference based methods are highly sensitive to heterogeneity is notably consistent.

One reason that BayesPrism performs well is that it explicitly accounts for heterogeneity in the malignant cells. In fact, it fits individual sub-types of malignant cells separately and reports the sum as the total malignant fraction. However, while this feature can be easily added to refernce based methods this did not recover BayesPrism performance levels (data not shown). On the other hand marker-based methods we investigate do not perform such explicit heterogeneity accounting, and are nevertheless still competitive with BayesPrism. This result indicates that the major advantage in robustness comes from the conceptual framework itself.

It may appear counter-intuitive that maker-based methods can outperform reference-based ones as marker-based methods seemingly use less of the available prior information. However, in the heterogeneous setting this turns out to be an advantage. Reference methods attempt to fit the true data generating process by solving a regression problem and differ in the choice of feature selection and regression objective. From this perspective of regression, the heterogeneous simulation introduces bias in the true mean and also induces a complex variance/covariance structure in the residuals which is not accounted for. On the other hand, marker based methods are highly robust to these effects as the residual variance and covariance of marker genes is low by construction and the exact mean values are not relevant.

In line with the view, BayesPrism presents an interesting case of a method that is fully quantitative but conceptually similar to the marker based approach. BayesPrism uses the full reference matrix but does so in a way that doesn’t rely on the exact reference values. One of the sampling steps of BayesPrism involves distributing the counts in the observed bulk expression for gene X over the current estimate of cell-type specific contributions with a multinomial distribution [40]. As such the absolute scale of gene X in the reference matrix is not relevant, as the values are interpreted as probabilities and normalized to sum to 1. Moreover, in this setting the contribution of a single gene to the final proportion estimate is directly proportional to its relative cell-type specificity times its expression value in the bulk sample. Consequently, the BayesPrism approach to a large degree negates the model misspecification sensitivity of regression-based methods as only the relative expression values for cell-type specific genes need to be correct. In this sense BayesPrism is actually a hybrid approach that lies somewhere between regression based and marker based methods. We expect that additional development in hybrid methods that combine the individual strengths of different approaches should be particularly effective at improving the performance in the realistic heterogeneous setting.

Our analysis provides valuable insights into the performance and tradeoffs of different conceptual approaches in a highly realistic simulation scenario thus establishing a framework for future methodological development. Beyond the specific deconvolution problem addressed in this work the heterogeneous simulation strategy can be employed in other simulation pipelines to produce more realistic performance benchmarks for additional tasks such as cell-type specific differential expression.

We also acknowledge some limitations of our approach. While we demonstrate that our heterogeneous simulation strategy matches the variance observed in real bulk samples, not all aspects of real data will be preserved. For example, the dependencies between cell types can be violated as we randomly combine cell-types from different patients. Chu et al.[15] found that certain biological pathway activation in malignant cells could be negatively correlated with cell type fractions of other non-malignant cells and the heterogeneous simulation we propose does not take into account such correlations. From the perspective of evaluating simulation quality these limitations could be detected as deviations from the expected covariance structure. Methods that overcome this limitation would need to take the ground truth cell-type covariance into account necessitating development of new proportion sampling strategies.

Reference-based approaches require prior information which can usually be obtained from single cell datasets that match the biological context or gene expression signatures of purified cell-types. However, as has been emphasized in multiple studies, in real practice, there could be technical variation between signature matrices and bulk mixtures due to differences in assay platform [31, 35]. Beyond technical differences arising from platform effects it is critical to consider biological differences between the reference/marker source and the dataset being analyzed. These are particularly important when legacy bulk datasets, that may come from rare or difficult to sample diseases, are analyzed using prior information derived from samples that are not matched in terms of cell composition and/or biological states of individual cell-types. In these cases, even enumerating the expected cell types may be a challenge. A complete evaluation for complex prior mis-specification will be the subject of future work.

Overall, our work suggests specific recommendations for creating realistic pseudobulk simulations and highlights counterintuitive findings regarding the performance of deconvolution approaches from different conceptual classes. Together, we expect that these contributions will provide the groundwork for future methodological improvements.

## Methods

### Single-cell RNA seq datasets and quality control

Four single-cell RNA sequencing datasets were used in this paper: (i) head and neck squamous cell carcinomas (HNSCC) tumors from Puram et al. [29], (ii) melanoma tumors from Schelker et al. [28], (iii) medulloblastoma (MB) tumors from Riemondy et al.[24] and (iv) another melanoma cohort from Jerby-Arnon et al.[27]. Data description can be found in Table 1. Single cell count data are CPM (count per million) normalized, where CPM values are divided by 10 to avoid inflation. We removed genes that are expressed in less than 5 cells and discarded genes from mitochondrial or ribosomal content. All expression matrices are in linear (non-log) space.

### Cell-type labels in scRNA datasets

For melanoma cohort 1 GSE72056, we used cell-type labels re-classified in Schelker et al’s study. For scMB dataset GSE155446, we re-annotate the immune population based on immune cell subtyping information from the interactive website of the original paper: https://d33sxa6bpqwi51.cloudfront.net/?ds=human%2FImmune-cells&meta=celltypes. We included major immune cell-types from their annotations for further study: DC, Neutrophil, NK cell, T cells. For all the macrophage subpopulations: chemokine myeloid, complement myeloid, M2-activated myeloid, non-activated microglia, we relabeled them into macrophages to ensure a reasonable resolution of cell-types. Immune cells that are classified as ‘Proliferate’ or do not have any subtyping label are excluded from further study. For the remaining scRNA datasets, we used their original cell-type labels.

### TCGA datasets

We downloaded the TPM normalized TCGA expression data from https://xenabrowser.net/ that has the same tumor type as the single cell data used in this study: GDC TCGA Head and Neck Cancer (HNSC) cohort and GDC TCGA Melanoma (SKCM) cohort. TCGA samples from primary tumor and metastatic tumors are selected for downstream variance comparison.

### Deconvolution benchmarking framework

For each scRNA dataset, the single cells are split into training (50%) and testing (50%) sets, where each set contains comparable numbers of cell-types to avoid unbalanced cellular compositions. Training cells are used to build reference matrices and cell-type specific markers, and testing cells are used to generate synthetic bulk samples. We then run deconvolution methods on the synthetic bulk samples and evaluate the performance by comparing the deconvolution results with the predefined cellular fractions. The above process is repeated 10 times for each scRNA dataset.

### Simulation of cell-type frequencies for synthetic bulk samples

To introduce variances into the cellular composition of synthetic bulk samples, we simulated cell-type frequencies that are close to that in real bulk samples. The cell-type proportions of each patient from the scRNA dataset were used as an approximation to the cell-type frequencies of real bulk samples. We fitted a beta distribution for each cell-type and drew random values from the fitted distribution as the simulated frequencies. Randomly selected frequencies for different cell-types are then scaled and summed to one for each synthetic bulk sample.

### Generation of synthetic bulk samples

Using testing cells from the data splitting step, three sets of bulk expression matrices containing 100 synthetic samples are generated based on the following strategies:

#### 1) homogeneous (random cells) simulation

We added up linear (non-log scale) expression values from n randomly selected testing cells and TPM (transcript per million) normalized the data, where n is the approximation of single cells contained in one bulk sample, (which is 500 for simulated HNSCC and melanoma samples, and 1500 for simulated MB samples), and the proportion of each cell-type corresponds to the simulated cell-type frequencies.

#### 2) semi-heterogeneous simulation

We restricted that the malignant parts of each synthetic bulk sample come from the same patient, while the non-malignant parts are randomly selected regardless of where they are from. Specifically, for each synthetic bulk sample *i,* the malignant expression signal come exclusively from a randomly selected patient’s malignant profile *C_malignant_* and is weighted according to the simulated malignant fraction; and the non-malignant single cells are randomly selected and weighted according to the corresponding simulated frequencies.

#### 3) heterogeneous simulation

We restricted that both malignant and non-malignant parts of each synthetic bulk sample come from the same patient. Specifically, for each synthetic bulk sample *i,* given a cell-type *k,* the expression signal of cell-type *k* comes exclusively from a randomly selected patient’s *k* profile C_k_ and is weighted according to the simulated fraction. Note that within one synthetic bulk sample, the malignant cells and cell-type k cells can belong to different patients.

### Calculation of biological variance in bulk samples

The following statistics are calculated as indicators of biological variance:

#### 1) coefficient of variation (CV)

for each gene *i* in the simulated and pseudobulk samples, we calculated CV values on the log transformed expression using the following formula:

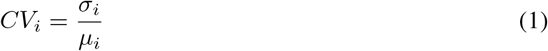

#### 2) average coefficient of variation (CV) for biological pathways

we downloaded the hallmark gene list from https://www.gsea-msigdb.org/gsea/msigdb/ and calculated the average CV values for genes included in each genelist, which is used as indicators for pathway variance.

#### 3) pairwise correlation between samples

for the simulated data, we first subset the expression to highly-variable genes (with *max.spec* greater than 0.5) and then calculated the Spearman correlation between each pair of simulated samples, with higher Spearman correlation suggesting lower variance between samples.

### Generation of reference matrices and cell-type specific markers for deconvolution

Using the training cells from the data splitting step, cell-type specific references are generated using four different approaches:

#### 1) CIBERSORTx signature matrix

we used the ‘Create Signature Matrix’ module from the online CIBERSORTx portal (https://CIBERSORTx.stanford.edu) to generate the signature matrix, with the training expression as input and all the parameters set to default values. We note that CIBERSORTx has a size limit for the input, so we downsampled the training cells and shrunk the input size when necessary. In the following approaches, reference matrices are built as a subset of reference profile C, where C is the averaged expression values for each gene across training cells from the same cell-type.

#### 2) autogeneS signature matrix

reference profile C is first filtered for highly variable (hv) genes and then used as autogeneS input. We used the following parameters in the autogeneS optimize function: ngen=3000, seed=0, mode = fixed, nfeatures=400. The resulting optimized reference matrix with pareto index 0 is selected as autogeneS signature matrix.

#### 3) DE gene-based signature matrix and cell-type markers using one-against-all statistics

R package limma is used to identify genes that are differentially expressed in each cell-type based on the log2 transformed CPM values. The contrast is set to one-against-all to find genes that are specific to one cell-type against all other cell-types combined. Genes with log fold change greater than 2 are considered as cell-type specific marker genes. To control the minimum number of markers in each cell-type, we introduced the parameter ‘*minimum_n_*’ that relaxes the log2FC threshold so that each cell-type will have appropriate numbers of markers. DE gene-based signature matrix is then built by filtering reference profile C for the marker genes.

#### 4) DE gene-based signatures matrix and cell-type markers using pairwise comparison statistics

get.exp.stat() function from R package BayesPrism is used to perform pairwise t-tests between every pair of cell-types. Genes with log fold change greater than 2 are considered as cell-type specific markers. To control the maximum number of markers in each cell-type, we introduced another parameter ‘*maximum_n_*’ to select the top n genes with the lowest p-values as cell-type markers. DE gene-based signature matrix is then built by filtering reference profile C for the marker genes.

### Highly variable (hv) genes

For computational efficiency, we selected highly variable genes as candidates to run autogeneS and DE analysis. We used the plot.scRNA.outlier() function from R package BayesPrism to calculate the maximum cell-type specificity score (for each gene. Genes with *max.spec* greater than a threshold value (0.5 for autogeneS and 0.3 for DE analysis) are selected for downstream analysis. This filtering narrows down the gene candidates from more than 10 thousands to thousands.

### Deconvolution methods

We applied the deconvolution methods on the linear scale (non-log) input, which is based on guidelines from previous benchmarking studies [16, 11]. A comprehensive evaluation of how data transformation could affect deconvolution results can be found elsewhere [11]. Three major categories of deconvolution methods are used in this study: reference-based, marker based and reference-free methods.

#### 1) Reference-free deconvolution

Linseed was run using R package linseed, we used the default parameters and set CellTypeNumber to the known value k for all the bulk mixtures. CAMfree was run using the CAM() function in R package CAMTHC with default parameters, and the candidate subpopulation number parameter was set to the known value k as well.

#### 2) Reference-based methods

##### i. Regression-based deconvolution

For our evaluation we applied the non-negative least squares (nnls) algorithm which uses squared error as the loss and non-negativity and proportion (sum is less than or equal to 1) constraints. In real practice, there are more reference-based algorithms available to choose from. For example, it is possible to use penalized (ridge, lasso and elastic net [41]) or robust regression [42]. The popular support-vector regression methods, CIBERSORT [43], can be viewed as regression with hinge loss. A detailed evaluation of fifteen different reference-based algorithms can be found in Avila Cobos et al. ‘s study [11]. Their study showed that nnls algorithm achieves good deconvolution performance while minimizing running time and required memory and that the reference matrix is more important than the specific formulation of the regression problem. Based on these observations, we applied nnls throughout our reference-based methods. Using the reference matrices C from the aforementioned step, we solved the deconvolution problem using the following formula: *M = CA*, where *M* is the expression values of the synthetic bulk mixtures, and *A* is the cellular composition matrix. The above regression problem was then solved by non-negative least squares (NNLS) regression.

##### ii. Marker-based deconvolution

Cell-type markers built from training expression matrices are used to guide marker-based deconvolution methods. We used the function AfromMarkers() from R package CAMTHC to estimate the actual cellular composition matrices. At the same time, we also calculated three extra unitless statistics for comparison: average expression, eigengene and gsva enrichment values.

##### iii. Bayesian method

The only Bayesian method evaluated in this study is BayesPrim. To create the ‘prism’ object required for BayesPrism, we input the expression profiles for all the training cells as reference, and set the cell.type.labels corresponding to the cell-types labeled in the original single-cell paper; for cell.state.labels, we used the same cell.type.labels for non-malignant cells (meaning that these cells have the same cell.type.labels and cell.state.labels), and annotated malignant cells with their source of origin as an indicator of their cell state. We set the update.gibbs=FALSE in the run.prism() function, meaning that we are using theta information from the first round of Gibbs sampling (initial theta). We also included a comparison where we set update.gibbs=TRUE and used theta information from the second round of Gibbs sampling (updated theta) in Fig S15, which shows no clear difference between using two kinds of theta. Since using initial theta has clear computational efficiency compared with updated theta, we applied the results from initial theta throughout our study.

In addition to the methods mentioned above, three additional deconvolution methods with built in reference or cell-type markers: epic, quantiseq and xCell were also implemented for cell-type composition estimation.

### Cell-type mapping

For deconvolution methods with built-in reference or cell-type markers, they predict cell-type fractions or scores that are different from the cell-types we used to generate the synthetic bulk samples. For example, xCell outputs 64 different immune cell scores, while our synthetic bulk samples contain only limited types of immune cells. To map the cell-types from these deconvolution methods to the corresponding cell-types used in our synthetic data, a manually annotated cell-type mapping dictionary was used to guide appropriate mapping between cell-types (SupplementaryTable2).

For reference-free methods without cell-type labels, we annotate each unlabeled cell-type by finding its maximally correlated cell-types fractions. Note that multiple unlabeled cell-types can be mapped to the same cell-type.

### Evaluation of deconvolution performance

Pearson correlation and root mean square error (RMES) values are used to evaluate the accuracy of different deconvolution methods. Specifically, for Pearson correlation, we focus on per cell-type correlation that compares between the estimated proportions with known compositions within each cell-type, with higher Pearson r corresponding to better performance. Average Pearson correlation values across cell-types are used as an overall correlation performance score. For RMSE values, we focus on a global comparison between the estimates and the real fractions for all cell-types altogether, with smaller RMSE values indicating lower absolute difference thus better performance. Variance in Pearson *r* and RMSE values across experimental repeats are used to evaluate the robustness and reproducibility of each deconvolution method.

### Single cell clustering

Visualization of single cell clustering was performed using tSNE in the R package Seurat based on the top 30 principal components of the top 8000 variable genes, with a perplexity value of 30.

### Pseudobulk samples from scRNA datasets

Pseudobulk samples are built by aggregating cells within the same patient: the CPM normalized count values of single cells are added up in linear (no log) space and then transcript per million (TPM) normalized.

### Cell-type markers from external resources

We collected immune cell markers from three different resources: Imsig [44], xCell [12] and scType [30]. Because of the large redundancy within different immune signatures (e.g for T cells, there are naive T, memory T and effector T cell signatures), we reorganized the marker list by finding the interactions between redundant gene lists and summarized our final marker sets in SupplementaryTable1.

## Availability of data and materials

Single-cell RNA sequencing data were obtained from Gene Expression Omnibus (GEO: https://www.ncbi.nlm.nih.gov/gds) under accession number listed in Table 1. Gene expression data from TCGA is downloaded from UCSC Xena Browser: https://xenabrowser.net/. Hallmark gene sets used in this paper are downloaded from The Molecular Signatures Database (MSigDB): https://www.gsea-msigdb.org/gsea/msigdb/. All codes are available at https://github.com/humengying0907/deconvBenchmarking.

## Acknowledgments

We acknowledge the TCGA Research Network for data generated in: https://www.cancer.gov/tcga. We acknowledge Casey Greene and Ariel Hippen for helpful discussion.

