## Supplementary Figures for "Heterogeneous pseudobulk simulation enables realistic benchmarking of cell-type deconvolution methods"

Fig S1. Patient-specific heterogeneity in malignant cells

Fig S2. Patient-specific heterogeneity in non-malignant cells from the HNSCC dataset

Fig S3. Patient-specific heterogeneity in non-malignant cells from melanoma cohort 1

Fig S4. Patient-specific heterogeneity in non-malignant cells from the MB dataset

Fig S5. Patient-specific heterogeneity in non-malignant cells from melanoma cohort 2

Fig S6. Variance comparison between simulation strategies

Fig S7. Variation of biological pathways between simulation strategies

Fig S8. Comparison between cell fraction simulation strategies

Fig S9. Deconvolution results on the simulated bulk samples generated from sc-HNSCC dataset

Fig S10. Deconvolution results on the simulated bulk samples generated from sc-melanoma cohort1

Fig S11. Deconvolution results on the simulated bulk samples generated from sc-MB dataset

Fig S12. Deconvolution results on the simulated bulk samples generated from sc-melanoma cohort2

Fig S13. Summarized evaluation of deconvolution results using global RMSE values

Fig S14. Average running time comparison for reference-based methods

Fig S15. Cell-type level performance comparison for BayesPrism using initial theta and updated theta in three different datasets

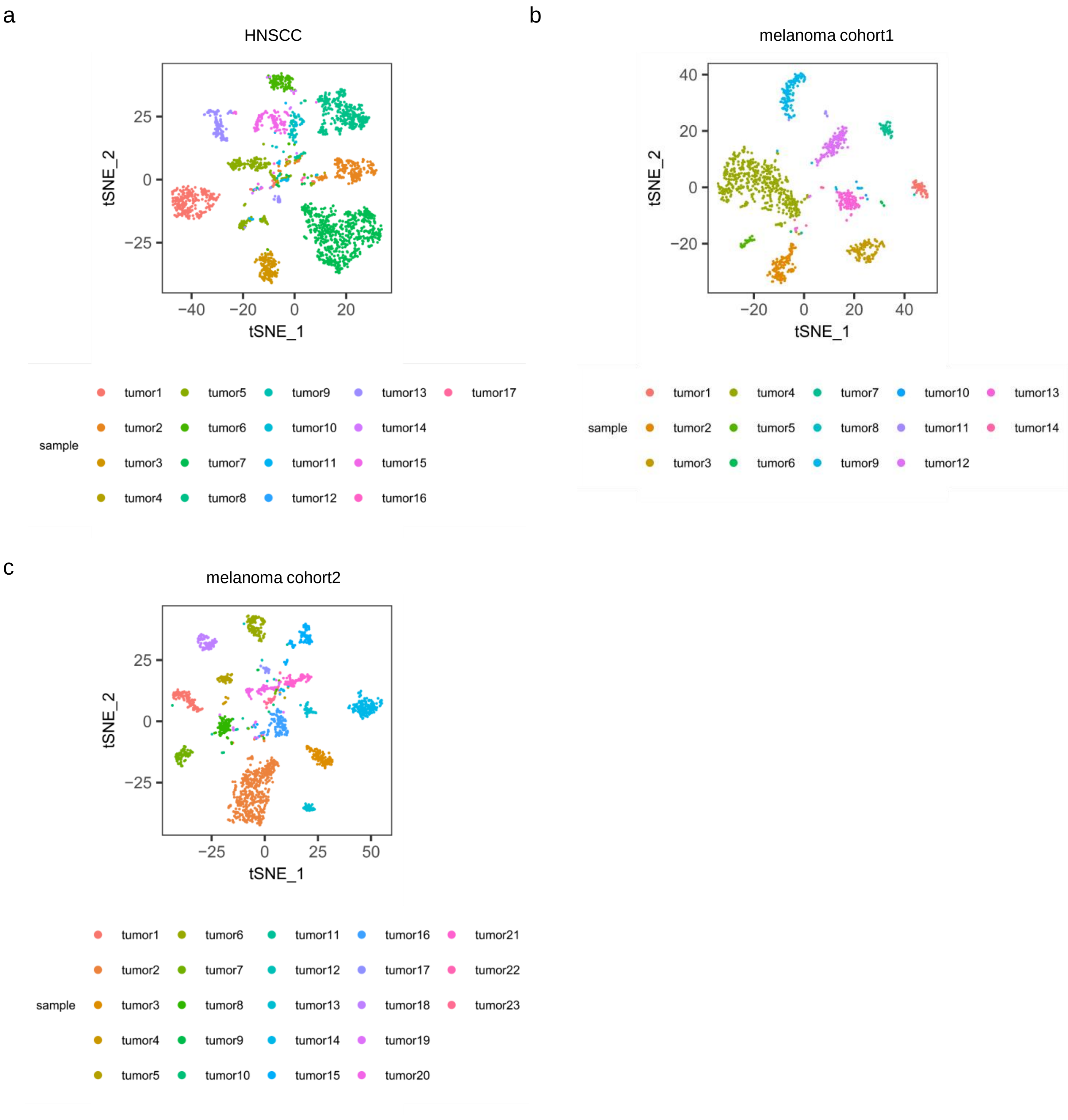


**Fig S1. Patient-specific heterogeneity in malignant cells**

(a-c) tSNE plot of malignant cells from single-cell HNSCC dataset (n=2,539), melanoma cohort 1 (n=1,310) and melanoma cohort 2 (n=2,018), colored by patient identifiers.

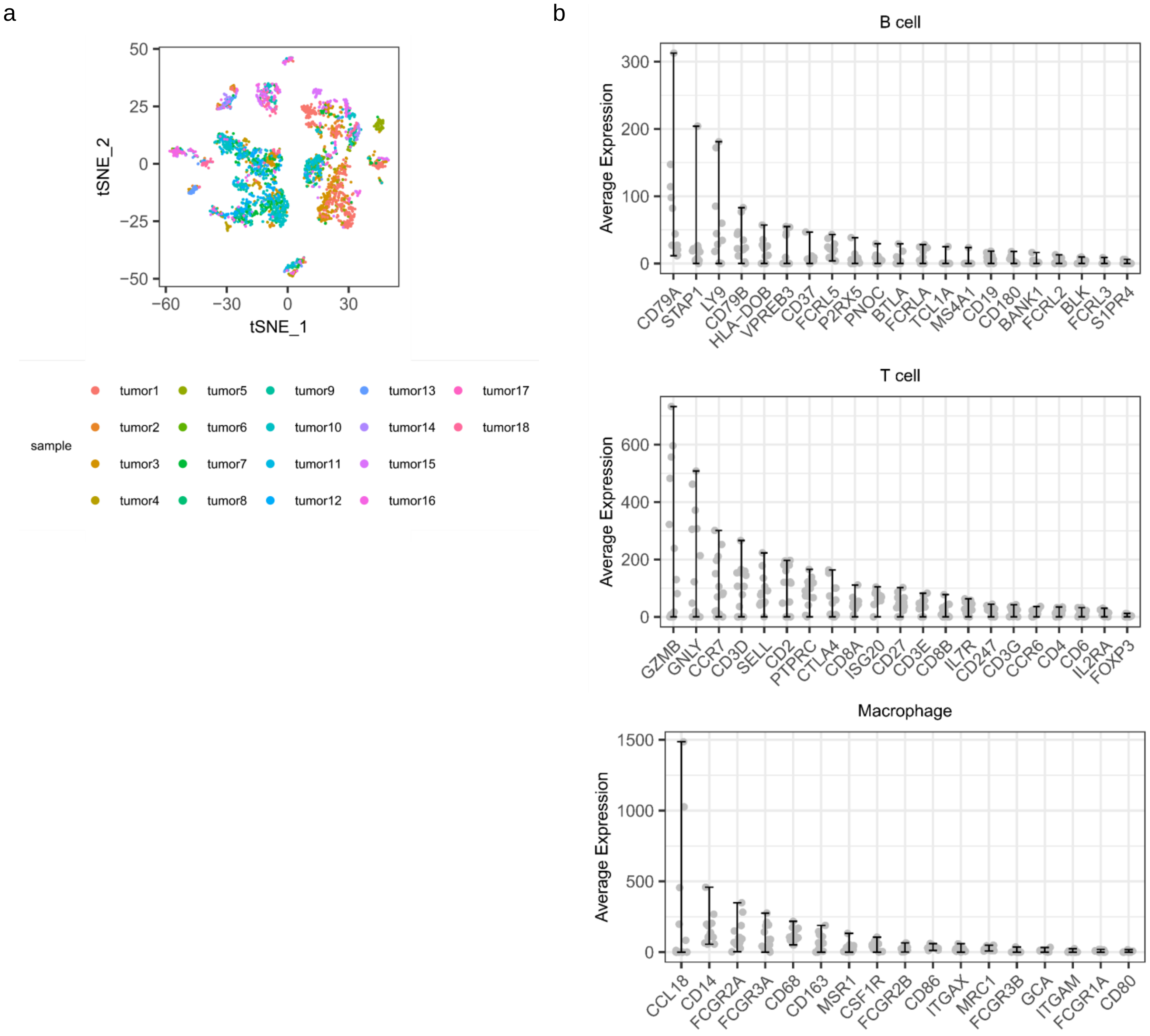


**Fig S2. Patient-specific heterogeneity in non-malignant cells from the HNSCC dataset**

(a) tSNE plot of all the non-malignant cells (n=2,539) from the sc-HNSCC dataset, colored by patient identifiers. (b) Average expression value of immune markers in each patient, which is aggregated over all the immune cells belonging to the same patient, with the error bar indicating the min and max average expression values.

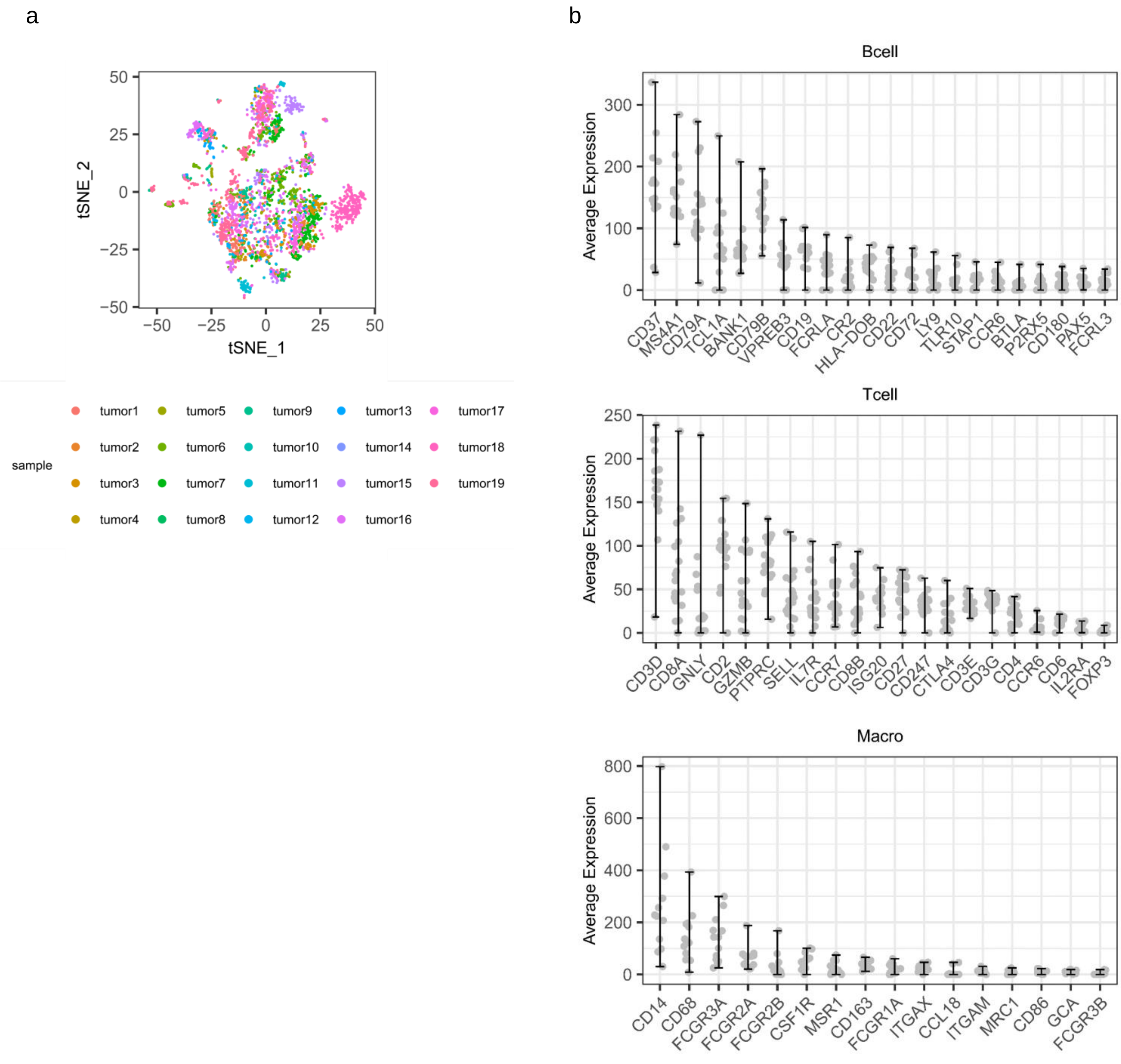


**Fig S3. Patient-specific heterogeneity in non-malignant cells from melanoma cohort 1**

(a) tSNE plot of all the non-malignant cells (n=1,310) from the melanoma cohort 1, colored by patient identifiers. (b) Average expression value of immune markers in each patient, which is aggregated over all the immune cells belonging to the same patient, with the error bar indicating the min and max average expression values.

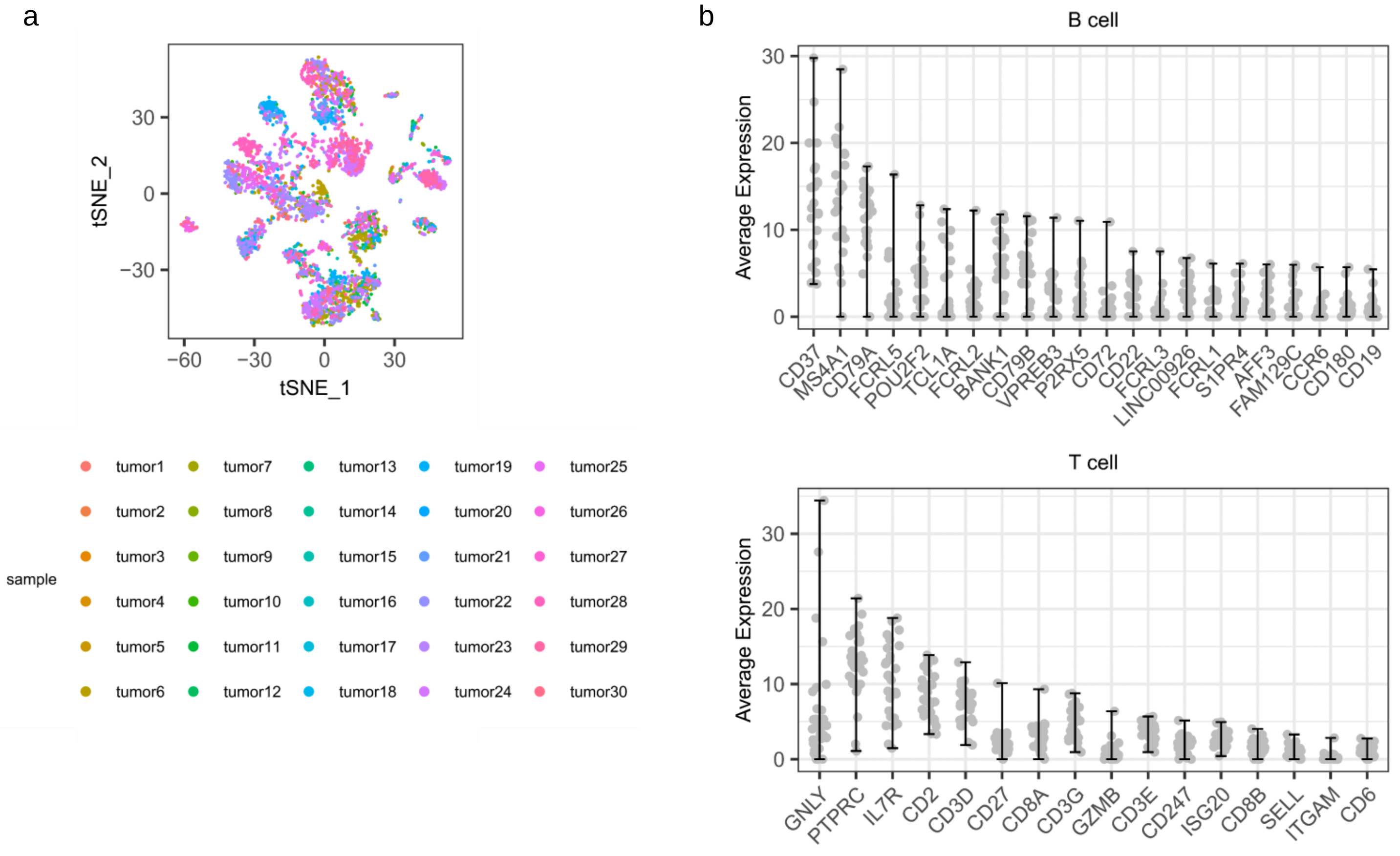


**Fig S4. Patient-specific heterogeneity in non-malignant cells from the MB dataset**

(a) tSNE plot of all the non-malignant cells (n=5,100) from the sc-MB dataset, colored by patient identifiers. (b) Average expression value of immune markers in each patient, which is aggregated over all the immune cells belonging to the same patient, with the error bar indicating the min and max average expression values.

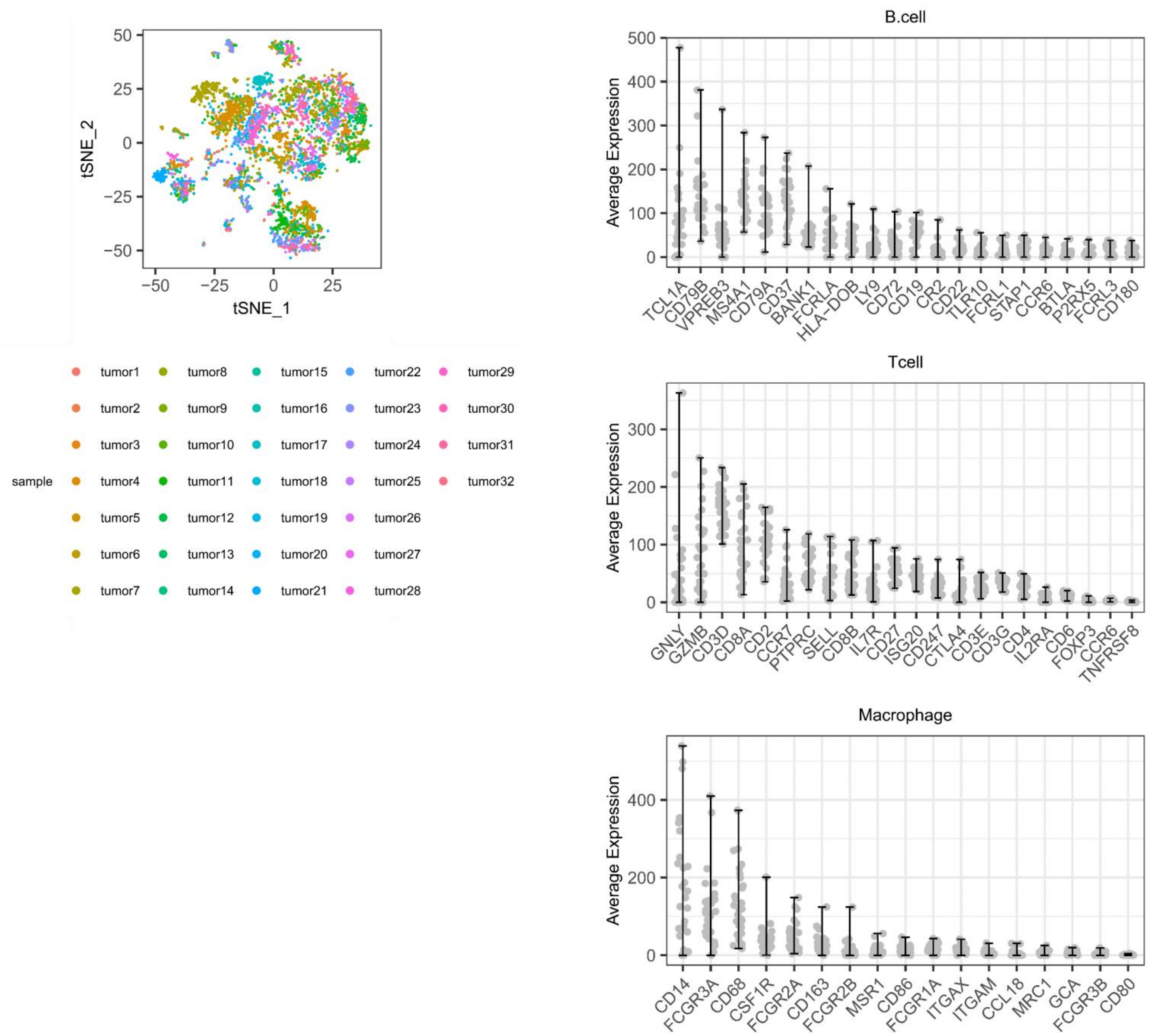


**Fig S5. Patient-specific heterogeneity in non-malignant cells from melanoma cohort 2**

(a) tSNE plot of all the non-malignant cells (n-4,816) from melanoma cohort 2, colored by patient identifiers.
(b) Average expression value of immune markers in each patient, which is aggregated over all the immune cells belonging to the same patient, with the error bar indicating the min and max average expression values.

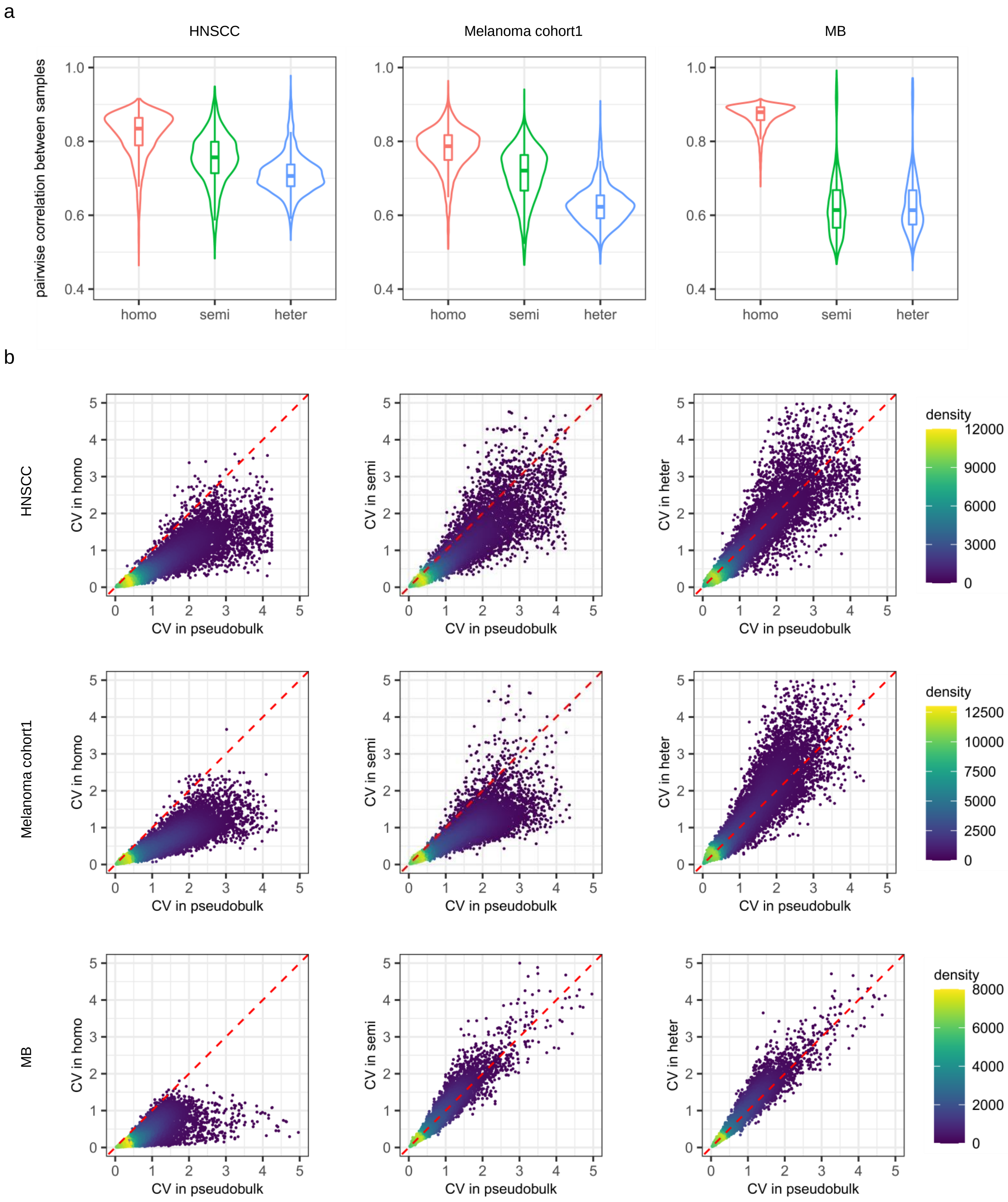


**Fig S6. Variance comparison between simulation strategies**

(a) Pairwise correlation between simulated bulk samples using homo, semi-heter, and heter simulation strategies from three sc-RNA datasets. (b) Scatter plots of coefficient of variation (CV) for all genes in the simulated bulk samples using homo, semi-heter, and heter simulation strategies from three sc-RNA datasets.

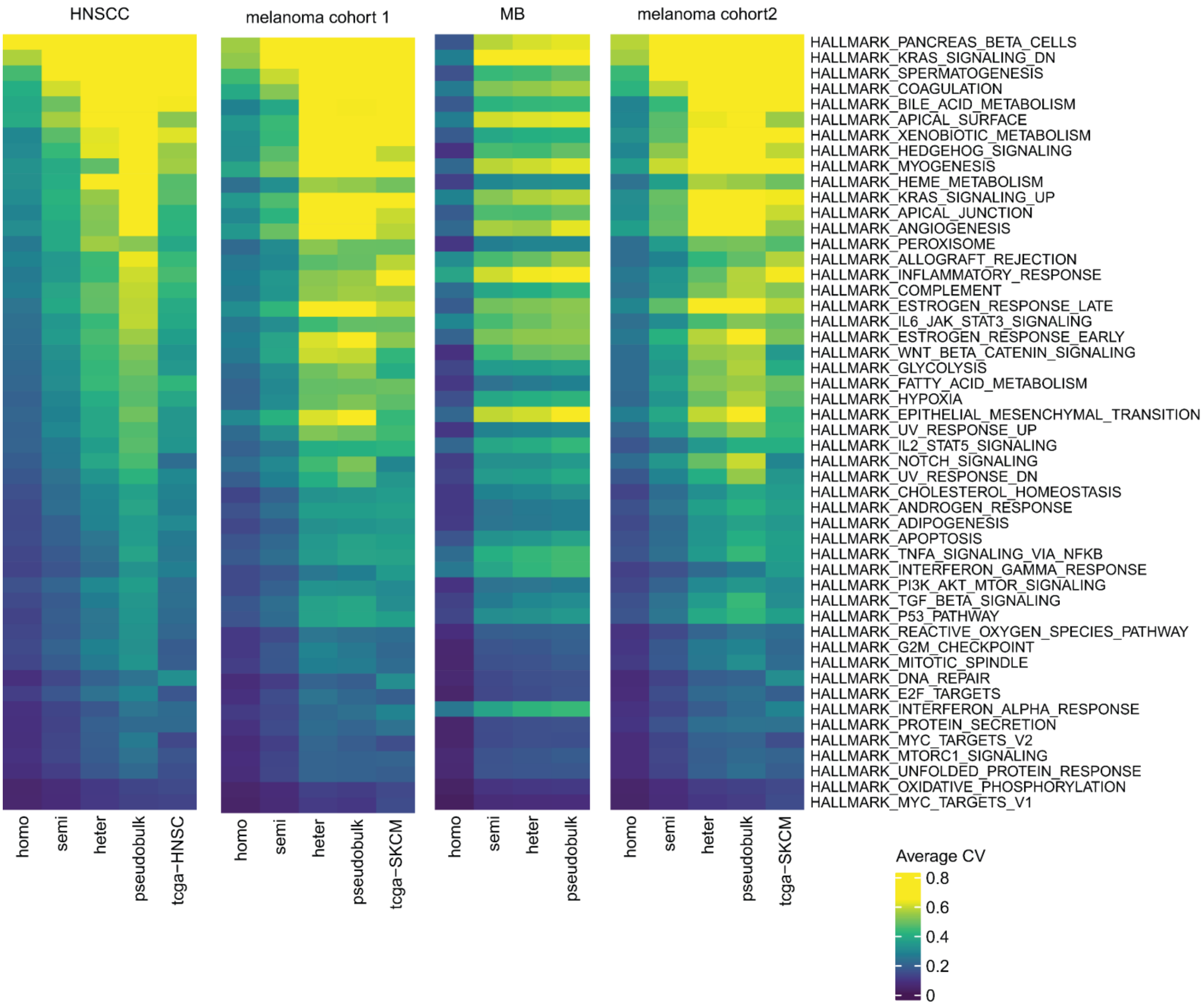


**Fig S7. Variation of biological pathways between simulation strategies**

Heatmap showing average coefficients of variance (average CV) for genes in 50 hallmark gene-sets in the simulated bulk samples, pseudobulk samples and real bulk samples from TCGA cohorts when paired tumor type is available.

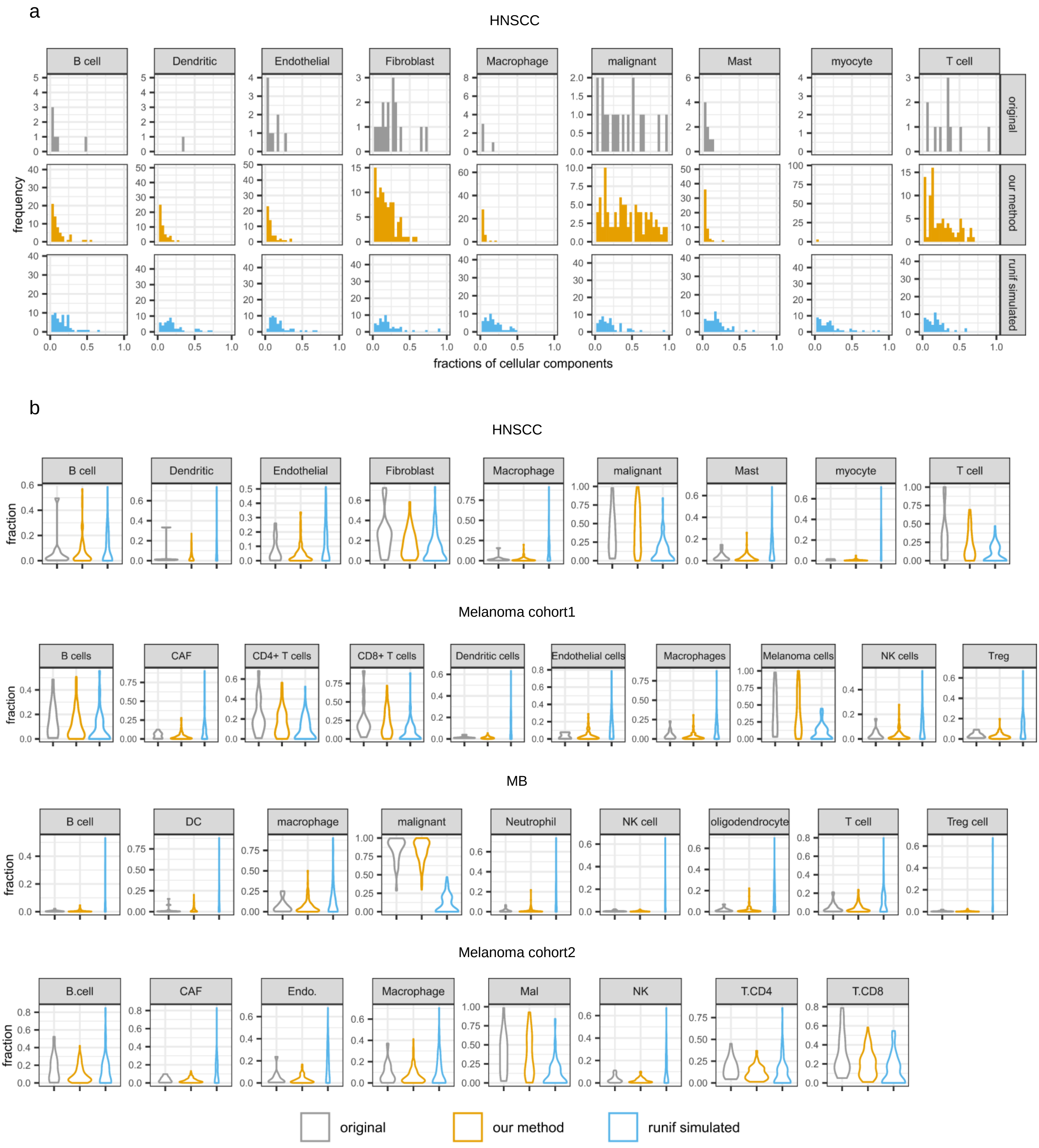


**Fig S8. comparison between cell fraction simulation strategies**

(a) Histogram showing the shape of proportion distribution with the first row: distribution of cell type fraction in 18 pseudobulk samples of the single-cell HNSCC datasets; second row: distribution of cell type fraction in 100 simulated bulk samples using our cell type fraction simulation strategy; third row: distribution of cell type fraction in 100 simulated bulks samples using runif simulation strategy. (b) Violin plot comparing cell-type proportion distributions in pseudobulk samples, simulated bulk samples using our simulation strategy and runif simulation strategy.

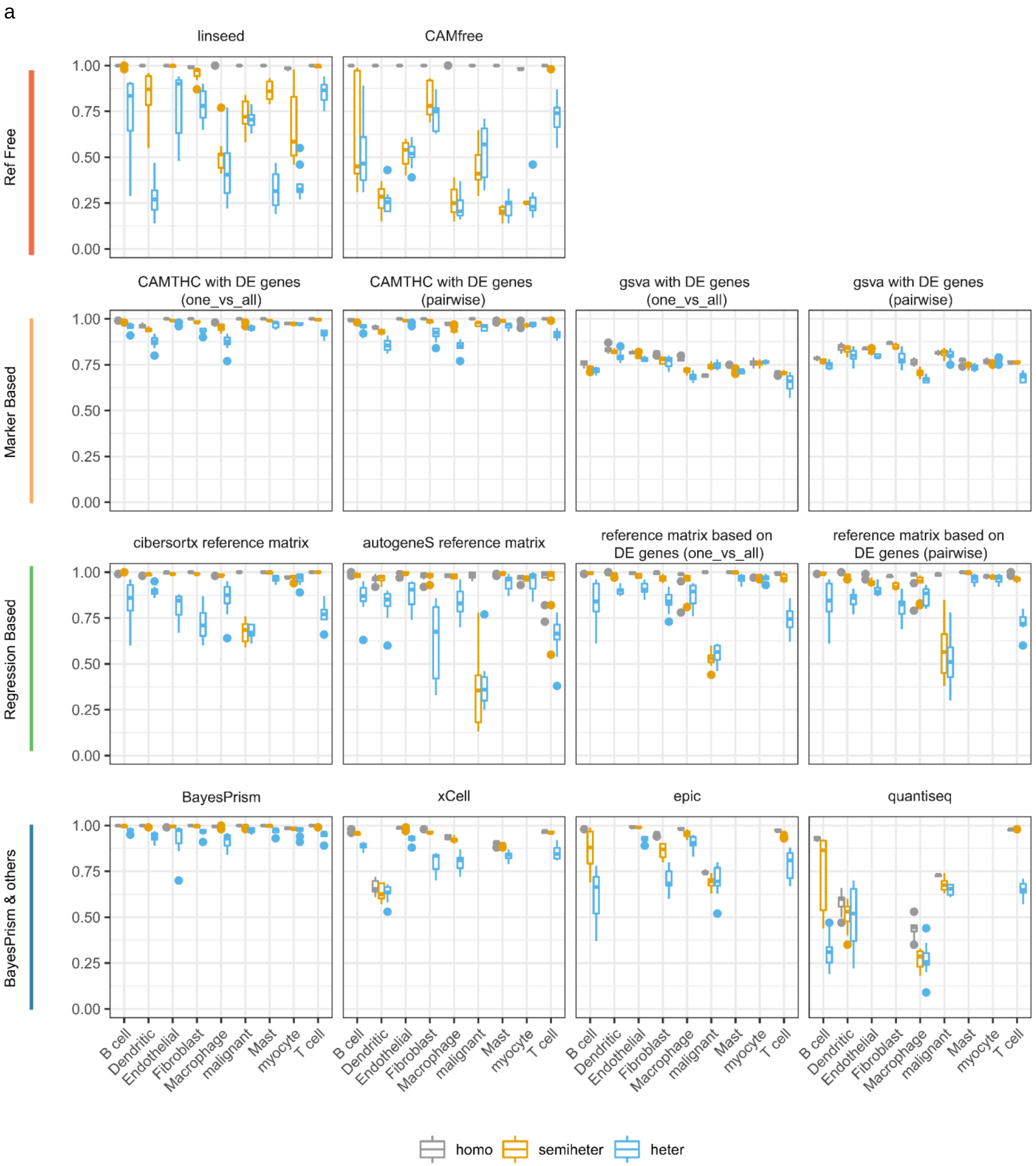


**Fig S9. Deconvolution results on the simulated bulk samples generated from sc-HNSCC dataset**

(a) Deconvolution performance, estimated by Pearson r (y axis) is summarized at per cell type level (x axis) for the simulated HNSCC bulk samples under different simulation strategy, with each box indicating the distribution of per cell-type level performance across 10 experimental repeats and colored by simulation strategies.

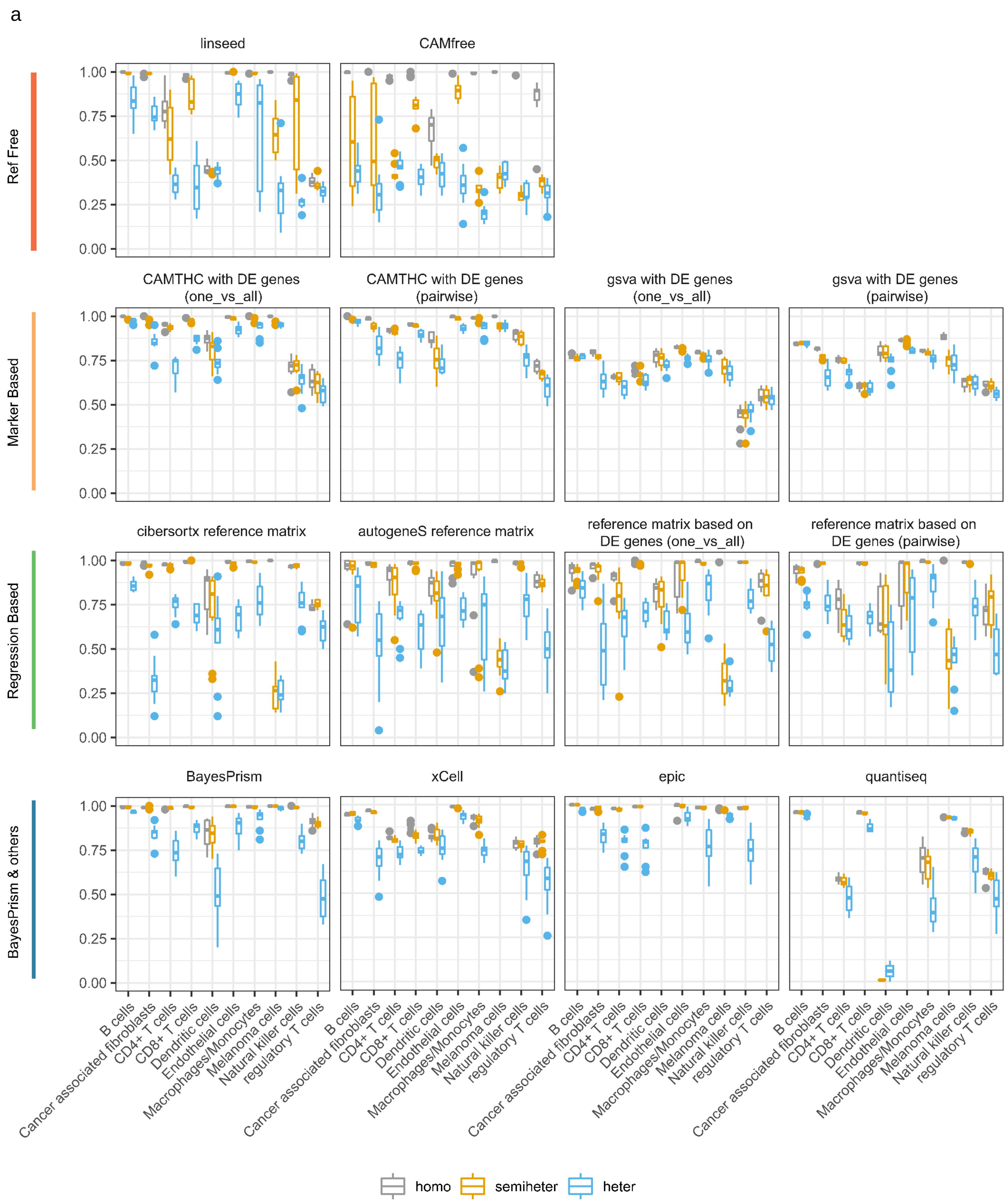


**Fig S10. Deconvolution results on the simulated bulk samples generated from melanoma cohort1**

(a) Deconvolution performance, estimated by Pearson r (y axis) is summarized at per cell type level (x axis) for the simulated melanoma bulk samples under different simulation strategy using single-cell melanoma cohort1, with each box indicating the distribution of per cell-type level performance across 10 experimental repeats and colored by simulation strategies.

a

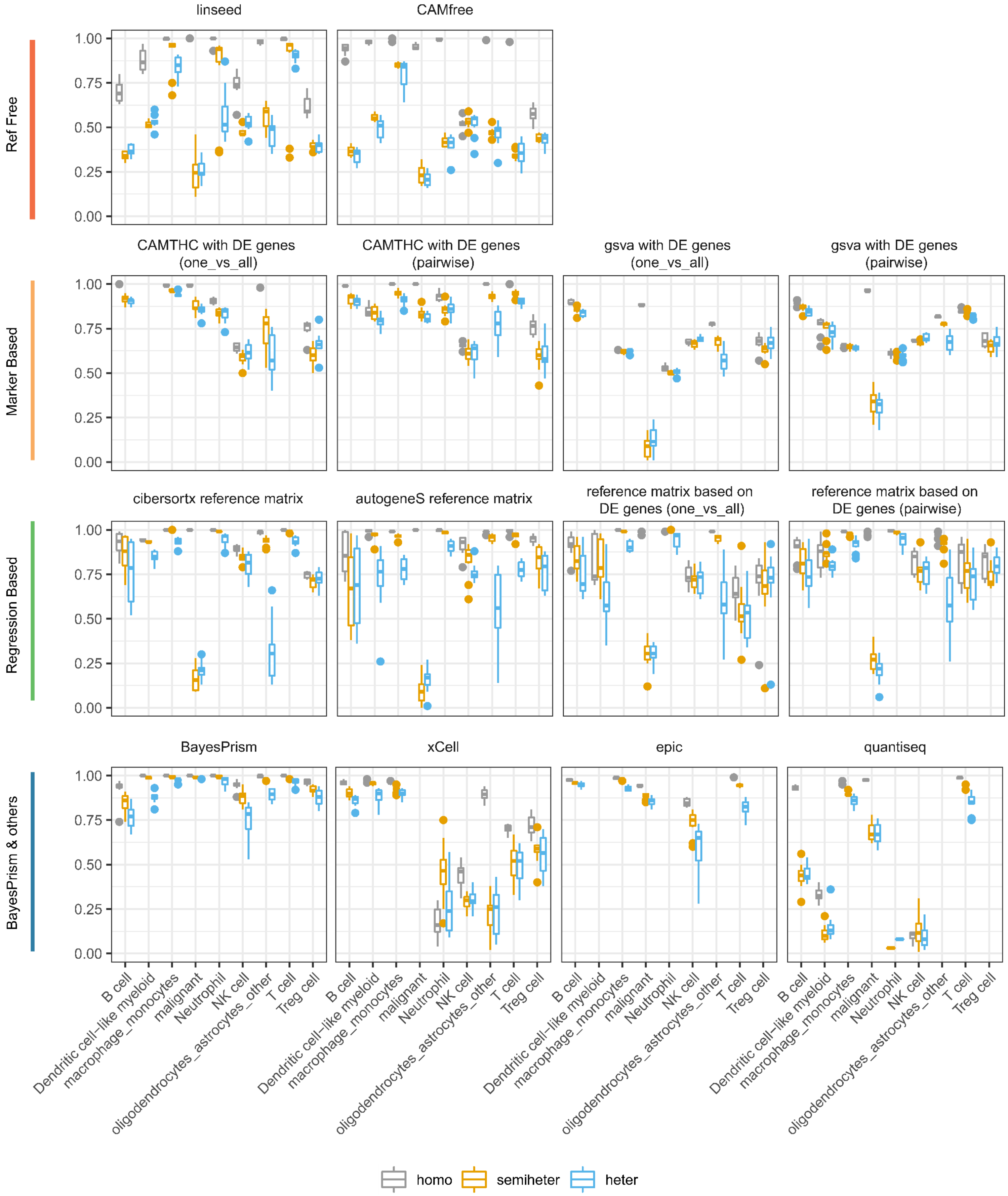


**Fig S11. Deconvolution results on the simulated bulk samples generated from sc-MB dataset**

(a) Deconvolution performance, estimated by Pearson r (y axis) is summarized at per cell type level (x axis) for the simulated melanoma bulk samples under different simulation strategy using single-cell MB dataset, with each box indicating the distribution of per cell-type level performance across 10 experimental repeats and colored by simulation strategies.

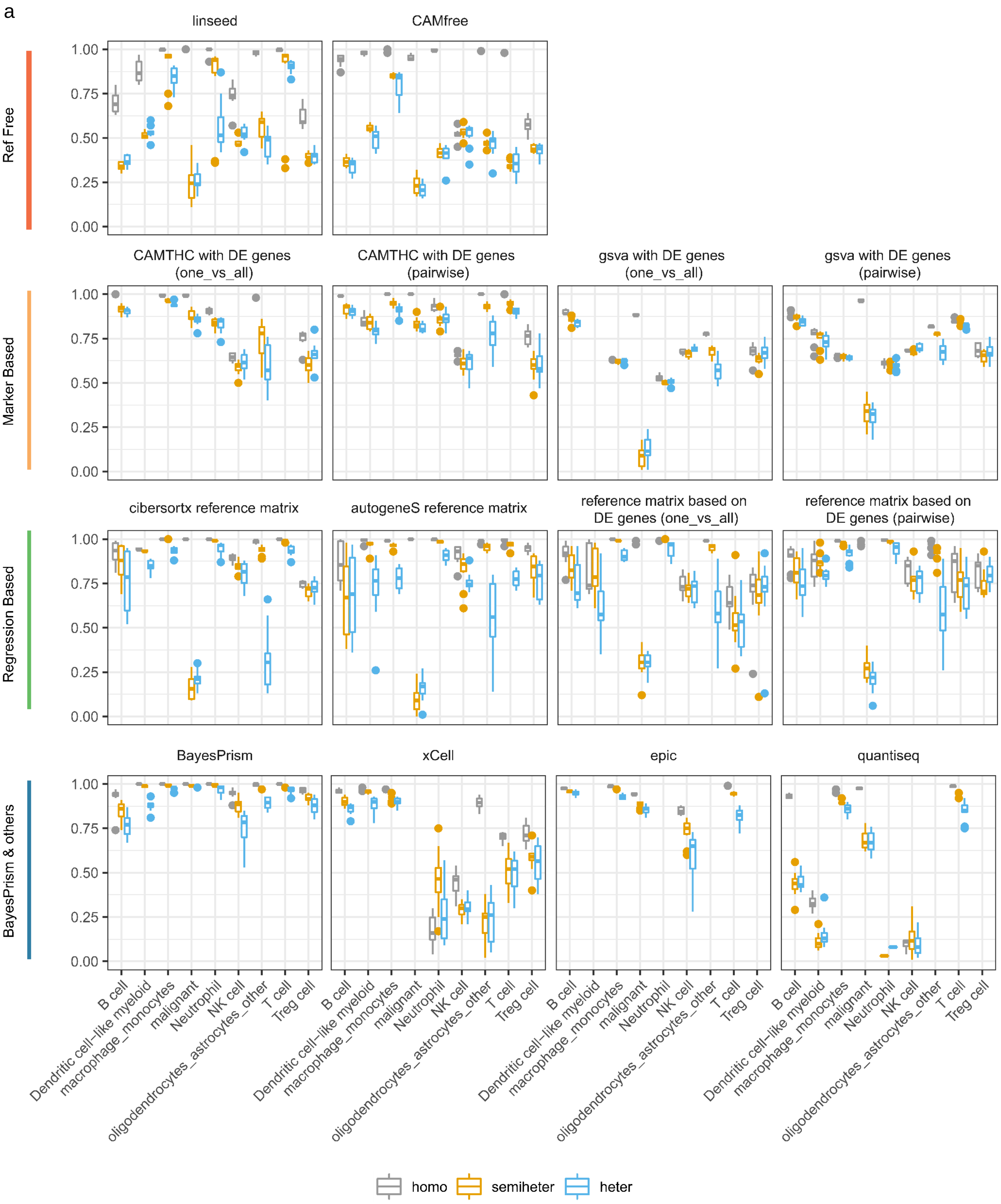


**Fig S12. Deconvolution results on the simulated bulk samples generated from sc-melanoma cohort2**

(a) Deconvolution performance, estimated by Pearson r (y axis) is summarized at per cell type level (x axis) for the simulated melanoma bulk samples under different simulation strategy using single-cell melanoma cohort2, with each box indicating the distribution of per cell-type level performance across 10 experimental repeats and colored by simulation strategies.

a

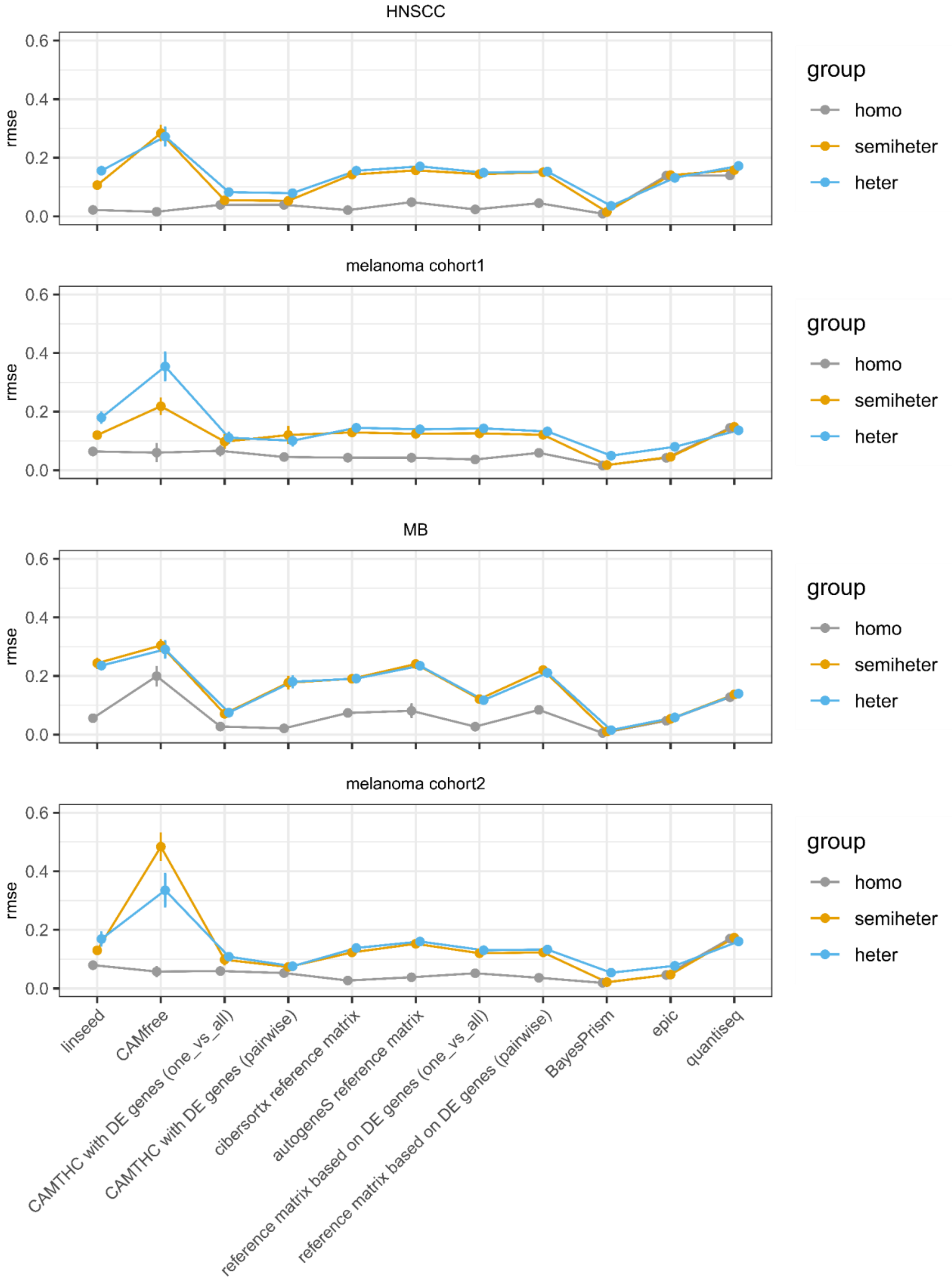


**Fig S13. Summarized evaluation of deconvolution results using global RMSE values**

(a) Lineplot showing global RMSE values of different deconvolution methods on bulk samples generated by different simulation strategies, with errorbar indicating the confidence intervals over experimental repeats.

a

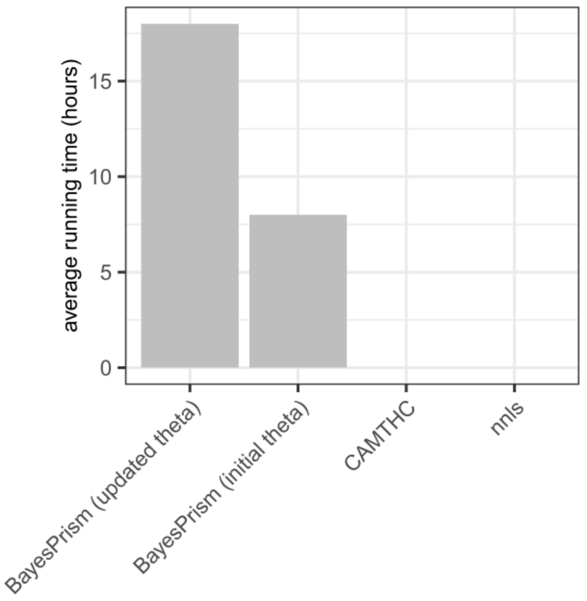


**Fig S14. Average running time comparison for reference-based methods**

(a) Boxplot comparing average running time required for reference-based methods: BayesPrism (with updated theta), BayesPrism (with initial theta), CAMTHC and nnls

a

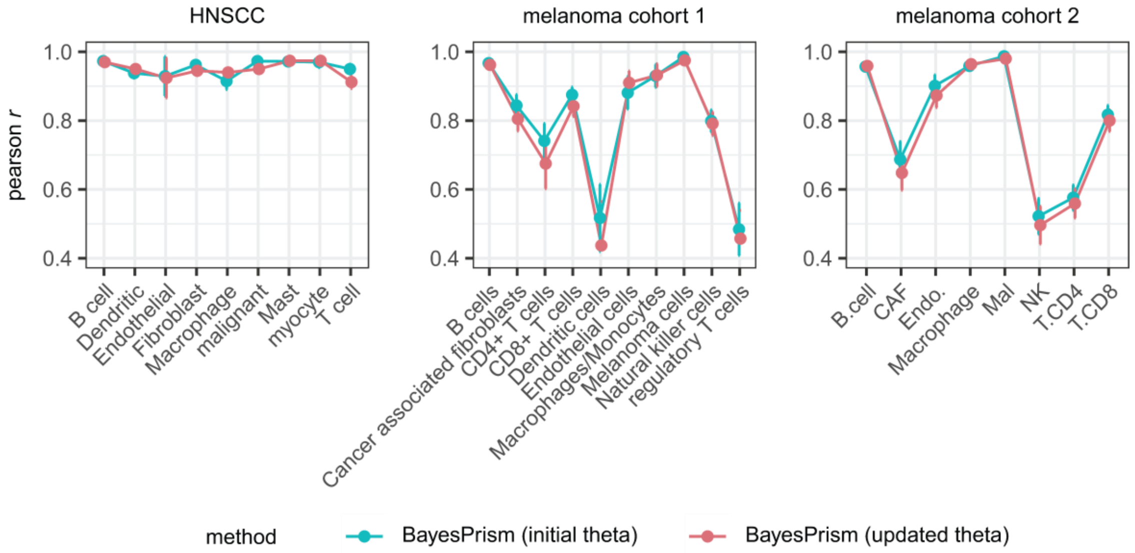


**Fig S15. Cell-type level performance comparison for BayesPrism using initial theta and updated theta in three different datasets**

(a) Lineplot comparing cell-type level performance when BayesPrism using initial theta (with update.gibbs=FALSE) and BayesPrism using updated theta (with update.gibbs=TRUE), with error bar indicating the confidence intervals over experimental repeats
